# Resilient anatomy and local microplasticity of naïve and stress hematopoiesis

**DOI:** 10.1101/2022.05.23.492315

**Authors:** Qingqing Wu, Jizhou Zhang, Courtney B. Johnson, Anastasiya Slaughter, Benjamin Weinhaus, Morgan McKnight, Bryan E. Sherman, Tzu-Yu Shao, Baobao Song, Marie Dominique Filippi, H. Leighton Grimes, Sing Sing Way, J. Matthew Kofron, Daniel Lucas

## Abstract

The bone marrow has an extraordinary capacity to adjust blood cell production to meet physiological demands in response to insults. The spatial organization of normal and stress responses is largely unknown due to the lack of methods to visualize most steps of blood production. Here we develop strategies to image multipotent hematopoiesis, megakaryopoiesis, erythropoiesis, and lymphopoiesis in mice. We combine these with imaging of myelopoiesis^1^ to define the anatomy of hematopoiesis in homeostasis, after acute insults, and during geriatric age. Blood production takes place via long-range migration of multipotent progenitors away from stem cells. Lineage-committed progenitors are then serially recruited to blood vessels where they contribute to lineage-specific microanatomical structures, composed of progenitors and immature cells, which function as production lines for mature blood cells. This anatomy is durable and resilient to insults as it was maintained after hemorrhage, acute bacterial infection, and with aging. Production lines enable hematopoietic plasticity as they differentially -and selectively-modulate their numbers and output in response to acute insults and then return to homeostasis. In geriatric mice the number of production lines is maintained, but their microanatomy becomes permanently remodeled in a cell- and lineage-specific manner indicating chronic anatomical adaptations to hematopoietic aging. Our studies uncover the sophisticated and durable anatomy of blood production -defined by distinct migratory behaviors depending on the maturation stage- and identify discrete microanatomical production lines that confer plasticity to hematopoiesis.

## MAIN

The spatial organization of cells in a tissue –its anatomy- dictates cell behavior and profoundly influences function^2^. Knowledge of the anatomy of each tissue is thus necessary to understand homeostasis and disease. Blood cell production takes place in the bone marrow through progressive differentiation of hematopoietic stem cells and progenitors. The bone marrow has extraordinary plasticity and quickly adjusts blood production to meet physiological demands in response to insults^3, 4^. Despite recent progress^5–10^ the anatomical organization of normal and stress hematopoiesis remains largely unknown. This is because current approaches do not allow simultaneous imaging of most types of hematopoietic progenitors and their daughter cell, in turn precluding in situ analyses of hematopoiesis; overcoming this hurdle is indispensable to defining parent and daughter cell relationships, changes in cell behavior during differentiation, and to identify the cells and structures enabling normal and stress hematopoiesis.

## RESULTS

Populations highly enriched in the different types of hematopoietic stem and progenitor cells (HSPC) have been defined through increasingly complex combinations of antibodies against defined cell surface markers^11–15^. Although scRNAseq analyses have revealed substantial heterogeneity of these populations^16^ antibody-based methods remain indispensable for identifying and isolating HSPC. Unfortunately, most of the antibody combinations used to prospectively isolate HSPC subsets by FACS are not suitable for confocal imaging analyses. We reasoned that unbiased analytical pipeline might reveal new combinations of surface markers to allow mapping of stepwise blood cell production in situ (Fig. 1a). We thus profiled 247 cell surface markers in phenotypically defined stem cells, multipotent progenitors, and lineage-committed myeloid and erythroid progenitors using three established cytometric strategies (Fig. 1a, Extended Data Fig. 1a, b and ^14, 15, 17^). To be useful for HSPC imaging a marker must: a) be expressed at sufficient levels for detection (which we experimentally determined to be an absolute fluorescence of 10^3^ over the background in our flow systems); and b) be able to discriminate between at least two types of HSPC. We thus selected markers that were uniformly expressed in at least one type of HPSC while being absent from one or more HSPC types. Thirty-five markers met these criteria (Fig. 1a, b).

**Figure 1.**
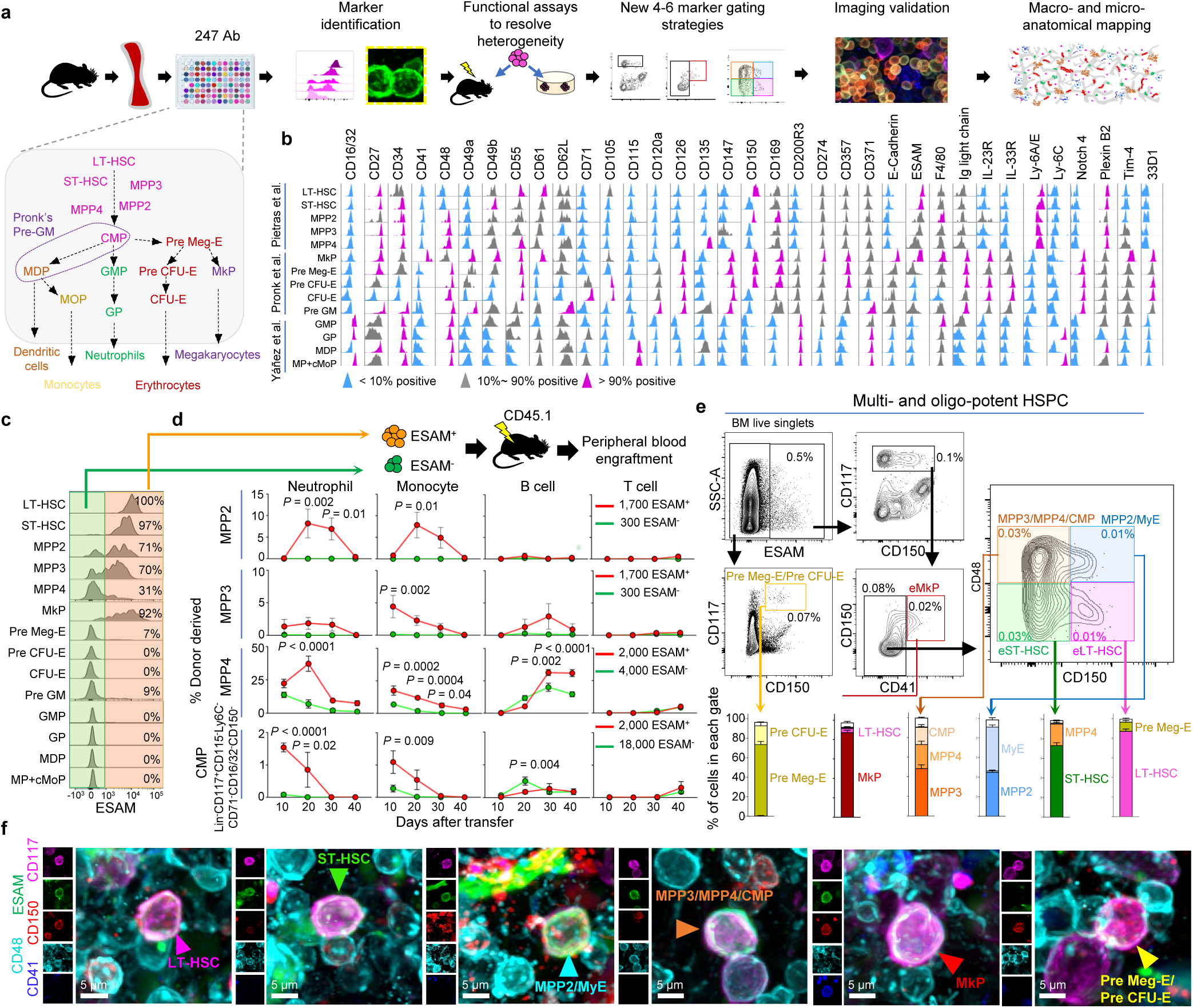
Identification of strategies to image hematopoiesis. **a,** Experimental pipeline. **b,** Histograms showing differential expression of 35 cell surface markers across 14 HSPC types. **c,** ESAM expression in indicated HSPC. **d,** Percentage of donor-derived cells in the peripheral blood after transplantation of the indicated progenitors (n = 5 mice for MPP2, 9 mice for MPP3, 4 mice for MPP4 and 4 mice for CMP per time point analyzed in 3 independent experiments). All data represent mean ± s.e.m. Statistical analysis was performed using Two-way ANOVA, followed by Sidak’s multiple comparisons test, adjusted *P* values are shown. **e,** FACS isolation strategy for indicated HSPC populations, the histograms show the percentage of classically defined progenitors in each gate (eLT-HSC = ESAM^+^ LT-HSC; eST-HSC = ESAM^+^ ST-HSC; eMkP = ESAM^+^ MkP. n = 5 mice in 3 independent experiments). All data represent mean ± s.e.m. **f,** Representative images showing identification of indicated HSPC in whole-mount sterna.

### ESAM enriches for –and allows imaging of– the most primitive stem and progenitor populations in the bone marrow

The immunophenotyping showed that ESAM is selectively expressed in 100% of LT-HSC, 97% of ST-HSC, 70% of MPP2 and MPP3, 31% of MPP4, 9% of Pre GM, 92% of MkP, and 7% of Pre Meg-E and absent in all other progenitors (Fig. 1b, c). To functionally resolve these heterogeneous populations, we performed transplantation experiments.

These revealed that all functional MPP2 and MPP3 are restricted to the ESAM positive fraction and that ESAM^+^ MPP4 are up to 5-fold more potent than ESAM^-^ MPP4 (Fig. 1d). Colony-forming assays revealed that the ESAM^+^ Pre Meg-E have monocyte and neutrophil differentiation potential whereas the ESAM^-^ Pre Meg-E have lost this capacity (Extended Data Fig. 1c). We thus defined ESAM^+^ Pre Meg-E as myeloerythroid (MyE) progenitors. Note that MyE are CD150^+^ and do not overlap with Pre GM which contain both CD115^-^ common myeloid progenitors (CMP) and CD115^+^ monocyte dendritic cell progenitors (MDP) (Extended Data Fig. 1a, d). While only ten percent of CD115^-^Ly6C^-^CD71^-^ CD16/32^-^CD150^-^Lin^-^CD117^+^ CMP express ESAM they contain all the engraftment potential in transplant assays indicating that bona-fide common myeloid progenitors are included in the ESAM^+^CD117^+^CD150^-^CD48^+^ population (Fig. 1c, d and Extended Data Fig. 1e). These findings demonstrate that ESAM identifies multi- and oligopotent progenitors in the bone marrow. They also led us to an isolation strategy replacing Sca1 and CD135 –which are classically used to prospectively isolate multipotent progenitors by FACS but do not work in immunofluorescence- with ESAM. This strategy allows simultaneous detection of highly purified LT-HSC, ST-HSC, megakaryocyte progenitors (MkP), MPP2/MyE (containing all functional MPP2 and MyE), and a mixed population containing the functional MPP3, CMP, and ESAM^+^ MPP4 (Extended Data Fig.1f). The eLT-HSC and eST-HSC gates are highly enriched in LT-HSC or ST-HSC (Fig. 1e) and have identical activity in competitive transplants as SLAM HSC (Extended Data Fig. 2a). Each of these six types of HSPC can be detected using five color immunofluorescences (Fig.1f) with similar frequencies when comparing imaging or FACS data indicating that the strategy detected all cells in the tissue (Extended Data Fig. 2b). ESAM also selectively stains blood vessels and megakaryocytes (Extended Data Fig. 2c) thus allowing simultaneous interrogation of these important components of the microenvironment.

### Imaging strategies for erythropoiesis and lymphopoiesis

Bona-fide CD150^+^ Pre Meg-E and and Pre CFU-E are uniformly ESAM^-^ whereas all other CD150^+^ HSPC are ESAM^+^ (Fig. 1c and Extended Data Fig.2d). CD71 -a classical marker of erythroblasts that is selectively upregulated in human CFU-E^18^- is highly expressed by all CFU-E, 70% of Pre CFU-E, and 30-80% of monocyte and neutrophil progenitors but absent in all other HSPC (Extended Data Fig.2d). To functionally resolve these heterogeneous populations we assessed erythroid differentiation potential using colony-forming assays. CD71^+^ Pre CFU-E contain almost all the erythroid activity whereas Ly6C^+^CD71^+^ progenitors were incapable of generating erythroid cells (Extended Data Fig. 2e). Expression analyses showed that mature hematopoietic cells do not coexpress CD71 and CD117 or CD150 and CD117 (Extended Data Fig. 2f). These led us to two FACS strategies to simultaneously detect all functional Pre Meg-E, Pre CFU-E, CFU-E, and ESAM^+^ HSPC or a mixed population of CD117^+^CD71^+^ Pre CFU-E and CFU- E, and classically defined early and late erythroblasts, reticulocytes, and erythrocytes by FACS and imaging (Extended Data Fig. 2g-k).

Common lymphoid progenitors (CLP) can be imaged as CD127^+^Lin^-^ cells^8^. All other steps of B cell maturation can be distinguished based on CD24, CD43, IgM, and IgD expression^19^. We combined these strategies to simultaneously image CLP, Pre-pro B, Pro B, and Pre B cells (Extended Data Fig. 2l, m).

Armed with these new imaging strategies we decided to examine the anatomy of hematopoiesis.

### Spatial segregation and long-distance migration of stem cells and multipotent progenitors

LT-HSC numbers and function are exquisitely regulated by adjacent niche cells^20^. A critical open question is whether LT-HSC and downstream progenitors colocalize –and therefore are regulated by the same niche ^6, 9, 21^- or, instead, progenitors leave the HSC niche as they differentiate. Imaging of 2-month-old mouse sternum segments showed that all stem cells and multipotent progenitors are found as single cells, evenly distributed through the bone marrow, and are no closer to each other than predicted from random simulations (median distances to closest progenitor > 100 μm (Fig. 2a-c and Extended Data Fig. 3a, b). This spatial segregation is not due to differential interaction with the microenvironment as most multipotent progenitors showed localization similar to that of LT-HSC: closer than random to megakaryocytes but farther than random from arterioles. Due to the abundance of sinusoids most multipotent HSPC localized within 10 µm of these vessels, but this distribution is no different from random cells (Extended Data Fig. 3c). These results indicate that multipotent progenitors move away from their parent and sister cells and migrate hundreds of micrometers as they differentiate. They also indicate that LT-HSC, ST-HSC, and other multipotent progenitors do not occupy the same niches.

**Figure 2.**
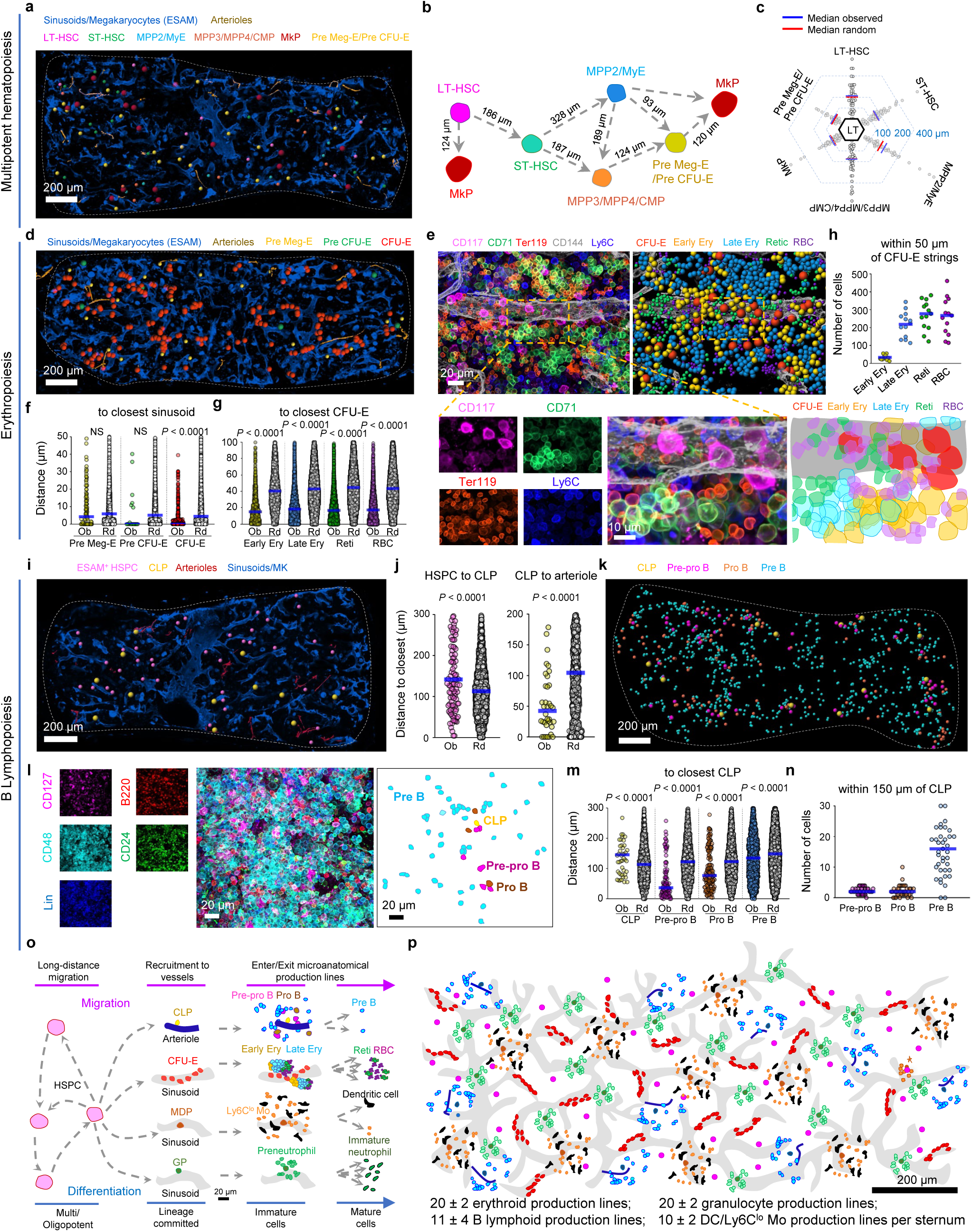
Anatomy of steady-state hematopoiesis in young mice. **a, b,** Map (a) and scheme summarizing (b) the distribution of the indicated HSPC in a 35-μm optical slice of the mouse sternum. Map dots are three times the average size of the relevant cell. **c,** Histogram showing the distribution of distances from each LT-HSC to the closest indicated cell (n = 41 LT-HSC in 5 sternum segments from 4 mice). **d,** Map showing the distribution of erythroid progenitors in mouse sternum. Map dots are three times the average size of the relevant cell. **e,** Maps and high-power image showing the distribution of late erythroid cells around CFU-E. Map dots are the average size of the relevant cell. **f,** Observed (Ob) and random (Rd) distance analyses from each erythroid progenitor to the closest sinusoid (n = 111 Pre Meg-E, 18 Pre CFU-E, and 627 CFU-E in 3 sternum segments from 3 mice). **g,** Distance analyses from each indicated cell to the closest CFU-E (n = 1,513 early erythroblasts (Early Ery), 6,506 late erythroblasts (Late Ery), 5,070 reticulocytes (Reti), and 5,388 red blood cells (RBC) in 5 sternum segments from 3 mice). **h,** Quantification of indicated cells within 50 µm of each CFU-E string (n = 13 CFU-E strings randomly selected in 3 sternum segments from 3 mice). **i,** Map showing the distribution of ESAM^+^ HSPC and CLP in mouse sternum. Map dots are three times (for ESAM^+^ HSPC) and five times (for CLP), the average size of the relevant cell. **j,** Distance analyses from ESAM^+^ HSPC to the closest CLP (left) and CLP to the closest arteriole (right) (n = 104 ESAM^+^ HSPC, 36 CLP in 3 sternum segments from 3 mice). **k, l,** Map and representative image showing the distribution of CLP, and B cell precursors in the sternum (the Lin panel contains CD2, CD3e, CD5, CD8, CD11b, Ter119, Ly6G, IgM, and IgD). Map dots are five times (for CLP) and three times (for Pre-pro B, Pro B, Pre B), the average size of the relevant cells. **m,** Distance analyses from each indicated cell to the closest CLP (n = 50 CLP, 104 Pre-pro B, 162 Pro B, 1,932 Pre B cells in 3 sternum segments from 3 mice). **n,** Quantification of indicated cells within 150 µm of each CLP (n = 41 CLP in 3 sternum segments from 3 mice). **o,** Scheme highlighting changes in migratory behavior during differentiation. **p,** Scheme showing the overall anatomy of hematopoiesis in 2-month-old mouse sternum. Statistical differences were calculated using two-tailed unpaired Student’s *t*-tests; *P* values are shown.

### Spatial organization of megakaryopoiesis

Megakaryocyte progenitors arise from LT-HSC, MPP2, or megakaryocyte erythroid progenitors^22^. Imaging analyses showed no specific spatial relationships between MkP and upstream progenitors, indicating that MkP also migrate away from their parent and sister cells (Fig. 2a-c and Extended Data Fig. 3b). In contrast, MkP selectively map near megakaryocytes (Extended Data Fig. 3c, d). To better understand megakaryocyte production we used *Ubc-creERT2:confetti* mice. In this model transient Cre activation leads to irreversible GFP, YFP, RFP, or CFP expression in 7.3% of total cells and 35.8% of megakaryocytes. This allows examination of clonal relationships in short-lived cells (Extended Data Fig. 3e and ^1^). We found numerous clusters of 2-4 sinusoidal megakaryocytes labeled with the same fluorescent protein –at much higher frequencies than predicted from random-indicating that they were produced by the same MkP. Unexpectedly, confetti-labeled MkP did not map near megakaryocytes labeled by the same color (Extended Data Fig. 3f, g). These results support a model where megakaryocyte progenitors are recruited towards active sites of megakaryopoiesis and terminally differentiate to generate 2-4 mature megakaryocytes.

### Erythropoiesis takes place in discrete production lines in the sinusoids

Pre Meg-E and Pre CFU-E are found as single cells through the tissue. Pre CFU-E migrate away from Pre Meg-E towards sinusoids (60% in direct contact with sinusoids) but do not map near CFU-E. These are found in large strings of 3 to 23 (Mean = 8 ± 4) CFU-E decorating the surface of a single sinusoid and away from arterioles (Fig. 2d-f and Extended Data Fig. 4a, b). Erythroblasts are selectively enriched near CFU-E but not Pre Meg-E or Pre CFU-E compared with random cells (Fig. 2g and Extended Data Fig. 4c). Indeed, all terminal erythroid cells were selectively enriched within 50 µm (the median distance for random cells) of a CFU-E (Fig. 2h). Higher powered images revealed that terminal erythropoiesis starts when CFU-E detach from the sinusoids, downregulate CD117 and upregulate CD71, progressively giving rise to several small clusters of early erythroblasts that bud from the vessel. These progressively upregulate Ter119 to generate large, nearly homogenous, clusters of 19 to 96 (Mean = 40 ± 4) late erythroblasts that, in turn differentiate into reticulocytes and erythrocytes that remain in close vicinity to the CFU-E strings (Fig. 2e). Confetti fate mapping showed that the CFU-E strings are oligoclonal while the erythroblast clusters are monoclonal (Extended Data Fig. 4d, e). These indicate that the CFU-E strings identify “erythroid production lines” in the sinusoids where CFU-E are serially recruited to generate defined numbers of red blood cells.

### B lymphopoiesis takes place in production lines near arterioles

We combined ESAM, CD117, and Lineage markers mixed with CD41 and Ly6C to map ESAM^+^ HSPC and CLP with arterioles and sinusoids (Fig. 2i and Extended Data Fig. 4f, g); CLP are found as single cells localizing far away (>150 µm) of multipotent HPSC (Fig. 2i, j). Arterioles are a niche for CLP^8^. In agreement, we found that CLP are selectively enriched near arterioles (mean distance = 53 µm) and depleted near sinusoids (Fig. 2i, j and Extended Data Fig. 4g). Most Pre-pro B, Pro B, and Pre B cells are selectively enriched near CLP, forming loose clusters (2 ± 1 Pre-pro B, 3 ± 2 Pro B, and 16 ± 8 Pre B within 150 µm of each CLP). The more mature cells located farther away from the CLP indicating migration away from the cluster (Fig. 2k-n). Unexpectedly, clonal fate mapping demonstrated that confetti-labeled CLP do not map near Pre-pro B, Pro B, or Pre B labeled with the same fluorescence protein (Extended Data Fig. 4h). These results indicate that the location of common lymphoid progenitors identifies oligoclonal B cell production lines near arterioles.

### Overall anatomy of hematopoiesis

We previously identified oligoclonal granulocyte production lines -where 1-2 granulocyte progenitors cluster with 10 ± 5 preneutrophils- and monocyte/dendritic cell production lines –where 1 MDP clusters with 15 ± 7 Ly6C^lo^ Mo and 6 ± 4 cDC- that selectively localize to distinct sinusoids (Extended Data Fig. 4i-k and ^1^). Neither granulocyte nor monocyte/dendritic production lines overlap with erythroid production lines (Extended Data Fig. 4l, m). Together the data reveals a sophisticated and complex organization of the bone marrow characterized by the long-distance migration of multipotent HSPC away from parent and sister cells. Lineage committed progenitors then migrate towards specific vessels where they are serially recruited to lineage-specific production lines with unique spatial and clonal architectures. Immature and mature cells leave these production lines to enter the circulation or localize to other bone marrow regions. The number of sterna production lines between mice is remarkably consistent with erythroid and granulocyte lines being the most abundant (in agreement with increased physiological demand for these two cell types, Fig. 2p, q and Extended Data Fig. 4n).

### The basic anatomy of hematopoiesis is maintained during -and after- acute stress

In response to acute insults the bone marrow initiates emergency differentiation programs leading to dramatic expansions and/or reductions in the output of one or more blood lineages. This is followed by a return to homeostasis once the insult is removed^1, 23–25^. The lack of tools to visualize differentiation has limited examination of these stress responses in situ. Key open questions are whether emergency blood production occurs via stress-specific anatomical structures or – alternatively- whether it exploits the existing structures present during homeostasis; whether these emergency responses are global (all structures in the tissue respond to the challenge) or local (only cells in discrete locations become perturbed); and whether the return to homeostasis also involves restoration of the preexisting anatomy. To explore these, we used models of acute hemorrhage – which causes expansion of erythropoiesis and reductions in lymphopoiesis- and acute infection with the intracellular bacterium *Listeria monocytogenes* -which causes expansion of multipotent progenitors and myeloid cells and reductions in erythropoiesis and lymphopoiesis (Fig. 3a and Extended Data Fig. 5a, b, and Fig. 6a and ^1^). Neither model showed major alterations in the vascular niches that sustain hematopoiesis (Extended Data Fig. 6b-d). Unexpectedly, despite major changes in blood production, the key anatomical features of hematopoiesis are maintained in both models of stress. We found spatial segregation and long-distance migration of multi- and oligo-potent HSPC (Fig. 3b and Extended Data Fig. 6e, note rare clusters of 2-4 MPP2/MyE after infection Fig. 3b and Extended Data Fig. 6f); lineage-committed progenitors selectively map to arterioles or sinusoids (except transient GP and MDP detachment in infection Fig. 3c, d and Extended Data Fig. 6g-n); and mature blood cells were generated in lineage-specific production lines (Fig. 3e and Extended Data Fig. 6o).

**Figure 3.**
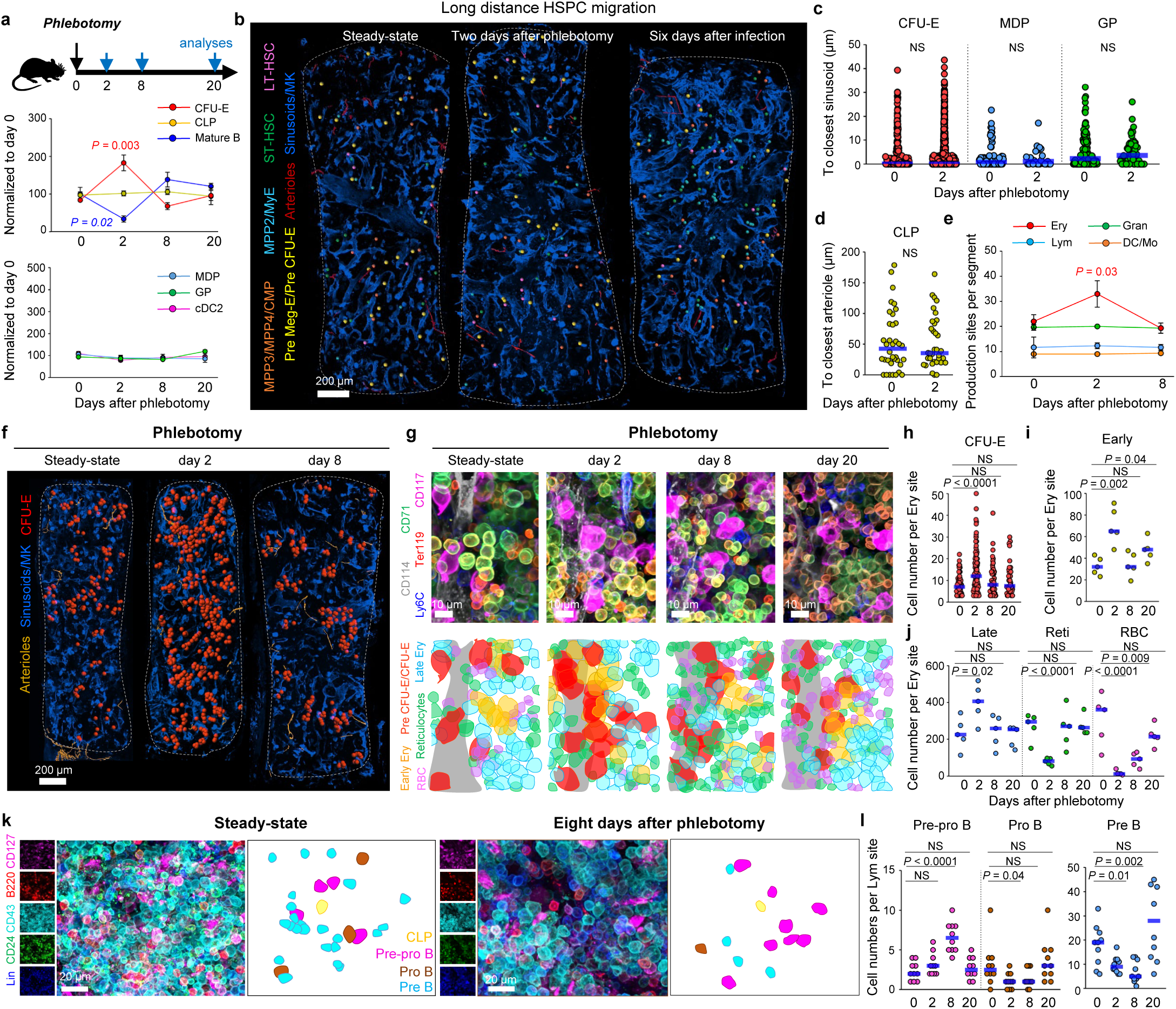
Anatomy of emergency hematopoiesis. **a,** Scheme showing the overall experiment design (top panel) to induce stress hematopoiesis and quantification (bottom panels) of changes in indicated HSPC by FACS (normalized to day 0) in response to phlebotomy (n = 4 mice). All data represent mean ± s.e.m. **b,** Maps showing minimal changes in HSPC distribution after phlebotomy or infection. Map dots are three times the average size of the relevant cell. **c,** Distance analyses from each indicated progenitor to the closest sinusoid. (n = 315 CFU-E, 62 MDP, 114 GP in 5 sternum segments from 5 mice in steady-states; n = 870 CFU-E, 25 MDP, 56 GP in 3 sternum segments from 3 mice 2 days after phlebotomy). **d,** Distance analyses from CLP to the closest arteriole (n = 36 CLP in 3 sternum segments from 3 mice in steady-states and n = 34 CLP in 3 sternum segments from 3 mice 2 days after phlebotomy). **e,** Quantification of each type of production line per sternum (n = 3 sternum segments from 3 mice for each time point). All data represent mean ± s.e.m. **f, g,** Maps (f) and representative images (g) showing the changes in erythroid production lines after phlebotomy. Map dots are three times the average size of the relevant cell. **h-j,** Number of CFU-E (h, n = 65, 99, 58, and 40 production lines for days 0, 2, 8, and 20, 3 sternum segments from 3 mice per time point) and indicated erythroid cells (i-j, n = 5 randomly selected production lines) in erythroid production lines after phlebotomy. **k, l,** Representative images (k) and number of cells per B cell production line (l) at the indicated time points after phlebotomy (n = 10 randomly selected production lines in 3 sternum segments from 3 mice for each time point). Statistical differences were calculated using two-tailed unpaired Student’s *t*-tests; *P* values are shown.

### Production lines orchestrate hematopoietic plasticity to stress

The data also showed dynamic adaptations of the production lines in response to stress. Two days after phlebotomy there are 50% more erythroid production lines, and these contain 75% more CFU-E but 1.5- fold fewer reticulocytes and 12- fold fewer erythrocytes as these are released into the circulation (Fig. 3e-j). Hemorrhage inhibits lymphopoiesis without reducing the number of B cell production lines (Fig. 3e and Extended Data Fig. 5a, 6g). Instead, there is a selective accumulation of Pre-pro B cells around the CLP but reductions in Pro B and Pre B cells indicating delayed differentiation at the Pre-pro B stage (Fig. 3k, l). These microanatomical changes are progressively restored until, on day 20 after phlebotomy, the anatomy of erythroid and lymphoid production lines is indistinguishable from steady-state mice (Fig. 3g-j, l). Phlebotomy did not affect the number and anatomy of neutrophil and monocyte/dendritic cell production sites at any time point examined (Fig. 3e and Extended Data Fig. 5a).

*L. monocytogenes* infection also causes lineage-specific microanatomical changes in the production lines. Reductions in lymphopoiesis take place via reduced output of B cell production lines, starting at the Pro B stage (Extended Data Fig. 7a, b), while the number of B cell lines is unaffected (Extended Data Fig. 6g, o). In contrast, reduced erythropoiesis occurs via reductions in both the number of erythroid production lines and their overall output (Extended Data Fig. 6o and Extended Data Fig. 7c- e). Infection causes minimal increases in the number of myeloid production lines (Extended Data Fig. 6o). However, it dramatically increases dendritic cell - and reduces Ly6C^lo^ monocyte- output in dendritic cell/monocyte production lines (Extended Data Fig. 7f-h). In agreement with our published studies infection also leads to increased GP numbers in the granulocyte production lines (Extended Data Fig. 6j, Extended Data Fig. 7i, j and ^1^). These perturbations are completely restored 20 days after the initial infection (Extended Data Fig. 7h, j).

These results demonstrate that the basic anatomy of hematopoiesis is durable and resilient to acute insults; that production lines orchestrate hematopoietic plasticity as they dynamically adapt their numbers and output to adjust blood production to demand –thus indicating that stress hematopoiesis uses the same structures as steady-state hematopoiesis for generating blood; that all production lines for a given lineage are synchronized as they simultaneously expand or contract in response to insults; that production lines for different lineages are independently regulated; and that the anatomy of hematopoiesis is fully restored once the acute insult is resolved.

### Chronic microanatomical adaptations to aging

Physiological aging represents a chronic insult that profoundly remodels the vascular niches that support hematopoiesis and leads to overall increases in cellularity, LT-HSC, and myeloid progenitor numbers (Extended Data Fig. 8a and^4, 26–28^). To examine how this chronic insult perturbs the anatomy of blood production we mapped hematopoiesis in 2- and 20-month-old mice.

A limitation of our imaging of multipotent hematopoiesis is that we use CD41 to distinguish MkP from other ESAM^+^ HSPC (Fig. 1e). However, aging causes accumulation of myeloid-biased CD41^+^ LT- HSC (Extended Data Fig. 9a and ^4^). This prevents using solely CD41 expression to discriminate between CD41^+^ MkP and CD41^+^ LT-HSC in aged mice. CD42d is exclusively expressed in MkP thus allowing imaging of bona-fide LT-HSC during aging (Extended Data Fig. 9a, b and^28^). Counterstaining with CD42d showed that CD41^+^ myeloid-biased and CD41^-^ lymphoid-biased LT-HSC are uniformly smaller than MkP (Extended Data Fig. 9c, d). This allowed us to use our original imaging strategy but now combining CD41 expression and cell size to simultaneously map CD41^+^ and CD41^-^ LT-HSC, MkP, and all other multi- and oligopotent progenitors with high sensitivity and specificity (Fig.4a and Extended Data Fig. 9e).

Imaging analyses confirmed profound remodeling of the vascular network that supports hematopoiesis. This is characterized by more numerous, shorter, and highly branched sinusoids and increased megakaryocytes in aged mice (Fig. 4a and Extended Data Fig. 9f-h). Unexpectedly, this barely perturbed the basic anatomy of hematopoiesis: we still found spatial segregation and long- distance migration of multipotent progenitors away from myeloid and lymphoid biased LT-HSC and each other (Fig. 4a, b and Extended Data Fig. 9i, j); selective localization of lineage-committed progenitors to vessels (except CLP, Extended Data Fig. 9k-m); and the number of production lines was maintained or slightly increased when compared to 2-month-old mice (Fig. 4c).

**Figure 4.**
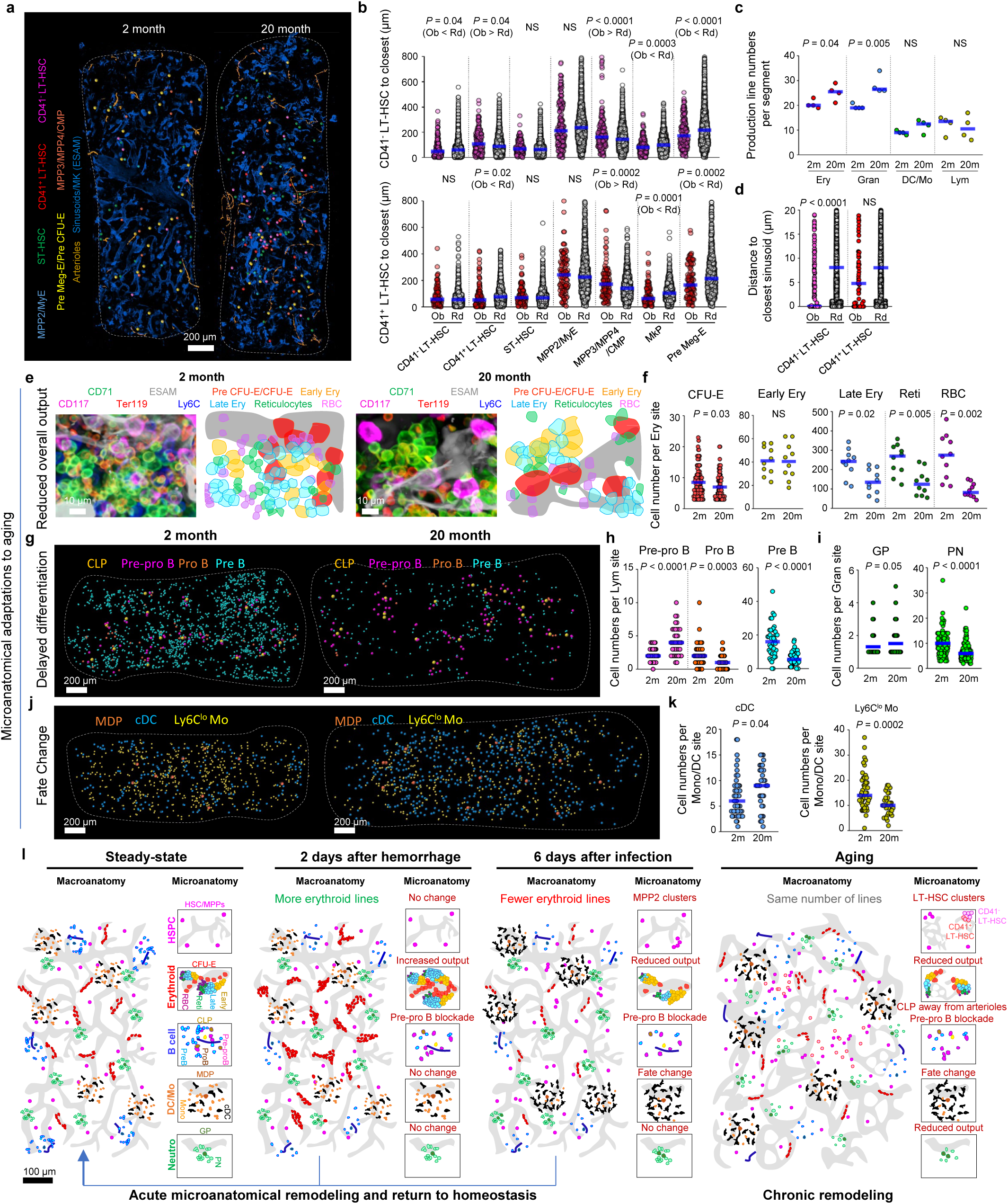
Anatomy of aged hematopoiesis. **a,** Maps showing changes in HSPC distribution with age. Map dots are three times the average size of the relevant cell. **b**, Distances analyses from each CD41^-^ or CD41^+^ (n = 191 or 109 in 3 sternum segments from 3 mice) LT-HSC to all other HSPC. **c,** Number of production lines per sternum segment 2- or 20-month-old mice (n = 4 sternum segments from 4 mice). **d,** CD41^-^ or CD41^+^ LT-HSC distance to sinusoids (*n* = 191 or 109 in 3 sternum segments from three 20-month-old mice). **e,** Representative images showing the changes in erythroid production lines in the aged sternum. **f,** Number of CFU-E (n = 96 and 88 production lines in 3 sternum segments from 3 mice) and indicated erythroid cells (n = 10 randomly selected production lines) in erythroid production lines in 2- and 20-month-old mice. **g, h**, Maps (g) and number of cells per B cell production line (h) in 2- (n = 41) and 20-month-old (n = 38 randomly selected B cell lines in 3 sternum segments from 3 mice). Map dots are five times (for CLP) and three times (for Pre-pro B, Pro B, Pre B), the average size of the relevant cells. **i.** Number of GP and preneutrophils (PN) cells per granulocyte production line in 2- (n = 96) and 20-month-old (n = 113 neutrophil production lines in 4-5 segments from 4 mice). **j, k,** Maps (n) and number of indicated cells (o) in 2- (n = 49) or 20-month-old (n = 34 DC/Ly6C^lo^ monocytes production lines in 3-5 segments from 3 mice). Map dots are three times (for MDP) and two times (for cDC and Ly6C^lo^ Mo), the average size of the relevant cell. **l.** Scheme summarizing microanatomical adaptations to stress. Statistical differences were calculated using two- tailed unpaired Student’s t-tests; *P* values are shown.

Strikingly, aging caused microanatomical perturbations that were cell- and lineage-specific: in young mice LT-HSC are found as single cells evenly distributed through the bone marrow (Fig. 2a-c). In contrast, in aged mice, lymphoid- and myeloid-biased LT-HSC localize much closer to other LT-HSC than predicted from random simulations –leading in some cases to the emergence of loose clusters of myeloid- and lymphoid-biased LT-HSC (Fig. 4a, b and Extended Data Fig. 9n). This correlates with selective lymphoid LT-HSC attachment to the remodeled sinusoids (Fig. 4d), which have been implicated in protecting HSC from aging^27^. The production lines for each lineage also showed lineage- specific microanatomical adaptations to age. The number of erythroid production lines increases slightly but each individual line shows reductions in almost all steps of erythropoiesis indicating reduced output (Fig. 4e-f). In other cases, we observe delayed differentiation: B cell lines display selective accumulation of Pre-pro B cells and severe reductions in Pro B and Pre B cells indicating stage-specific delayed differentiation at the Pre-pro B stage (Fig. 4g, h and Extended Data Fig. 10a). Granulocyte production lines also showed delayed differentiation (Fig. 4i and Extended Data Fig. 10b, c). Finally, monocyte and dendritic cell production lines switched from preferential monocyte production in young mice to preferential dendritic cell production in old mice (Fig. 4j, k and Extended Data Fig. 10d). Together with the data for acute stress (Fig. 3) the results also demonstrate that lineage-specific remodeling of microanatomical production lines enable hematopoiesis plasticity in response to acute insults and aging (Fig. 4l).

## Discussion

The bone marrow was the last major tissue where the spatial relationships between progenitors and differentiating cells – the anatomy- remained uncertain. This was due to a lack of validated cell surface markers to image the different types of hematopoietic progenitors. To overcome this problem, we generated databases of curated cell surface markers –and validation pipelines- that can be combined to prospectively image populations highly enriched in any hematopoietic progenitors of interest. We used these to develop strategies to map stepwise blood cell production, and defined the basic anatomy of hematopoiesis in the steady-state and how it adapts to acute insults and old age. We discover a sophisticated and elegant anatomy of hematopoiesis. It is characterized by long- distance migration of multi- and oligopotent progenitors; recruitment of lineage-committed progenitors to distinct blood vessels; and defined microanatomical production lines responsible for producing mature cells for each major blood lineage. This basic anatomy is durable, resilient to acute insults, and maintained through the adult lifespan. Finally, we show that the bone marrow adjusts blood cell production to respond to insults through microanatomical plasticity of the production lines.

Hematopoietic stem cells migrate through the bone marrow^10^ to the blood, circulate through tissues, and then drain into the lymph^29^. Our data indicates that this migratory behavior is also intrinsic to all steps of blood production. However, hematopoietic progenitors exhibit distinct migratory behaviors depending on maturation stage and lineage decisions. Whereas stem cells and multipotent progenitors migrate away from each other over long distances without obvious directionality committed progenitors selectively migrate towards lineage-specific production lines in –or near- blood vessels. CSF1^+^ sinusoidal endothelial cells and Oln^+^ periarteriolar cells represent discrete niches that recruit and regulate monocyte dendritic cell progenitors and common lymphoid progenitors respectively^1, 8^. We anticipate that production lines for other blood lineages will also be organized by specialized stromal and endothelial cells. Defining the identity of these local microenvironments and their role in regulating progenitor migration and differentiation in homeostasis and stress will be the focus of future studies. We propose that these production lines and local microenvironments will persist through life. However, a limitation of our study is that we cannot rule out that these lines are motile -or transient- as we cannot track the same marrow over time. We also have not explored other particularly interesting conditions that profoundly perturb the bone marrow like leukemia and bone marrow transplantation. The work shown here however provides the field with the tools and knowledge necessary to study stepwise blood production in situ in these and other physiological and pathological conditions.

## Extended Data Figure Legends

**Extended Data Figure 1. Resolving functional heterogeneity of HSPC subsets. a, b,** FACS plots showing the gating strategy to identify the 14 indicated hematopoietic progenitors (LT-HSC, ST-HSC, MPP2, MPP3 and MPP4 are identified as described by Pietras et al.^17^; MkP, Pre Meg-E, Pre CFU-E, CFU-E and Pre GM are identified as described by Pronk et al.^14^; GMP,GP, MDP and MP+cMOP are identified as described by Yanez, A. et al.^15^) interrogated through immunophenotyping. **c,** FACS plots and colony-forming assays showing that Lin^-^CD117^+^Sca1^-^CD41^-^CD16/32^-^CD105^-^CD150^+^ Pre Meg-E from (a) are heterogeneous and contain ESAM^+^ cells with myeloid and erythroid colony-forming activity (ESAM^+^ MyE) and bona fide Pre Meg-E (ESAM^-^) (n = 6 from 3 independent experiments). All data represent individual values with median plot. Statistical differences were calculated using multiple two-tailed unpaired Student’s t-test; *P* values are shown. **d,** FACS plots showing that the Lin^-^CD117^+^Sca1^-^ CD41^-^CD16/32^-^CD105^-^CD150^-^ Pre GM from (a) are heterogeneous and contain both common myeloid progenitors (CMP) and monocyte dendritic cell progenitors (MDP). **e,** FACS plots showing the gating strategy to identify pure CMP and ESAM and CD48 expression in these cells. **f,** FACS plots showing the relative purity of the HSPC identified using the strategy shown in Fig.1e. (eLT-HSC = ESAM^+^ LT- HSC; eST-HSC = ESAM^+^ ST-HSC; eMkP = ESAM^+^ MkP).

**Extended Data Figure 2. Validation of ESAM-based strategies to examine multipotent HSPC, erythropoiesis and B lymphopoiesis. a,** Overall experiment design (Top panels) and percentage of donor-derived cells (bottom panels) in lethally irradiated CD45.1^+^ recipients transplanted with the indicated amount of CD45.2^+^ SLAM (isolated as in Extended Data Fig. 1a) and ESAM (isolated as in Fig. 1e) LT-HSC (left) and ST-HSC (right) and 10^6^ CD45.1^+^ total BM competitor cells (n = 7 recipient mice from 3 independent experiments per time point). Statistical analysis was performed using Two-way ANOVA, followed by Sidak’s multiple comparisons test. It shows no significant difference for reconstitution ability between ESAM LT-HSC and SLAM LT-HSC, or ESAM ST-HSC and SLAM ST-HSC. **b,** Cell frequencies detected by FACS (white) or confocal imaging (orange) when using the isolation strategy shown in Fig. 1e (n = 9 mice for each group). **c,** Representative images and quantification showing that ESAM correctly labels all CD31/CD144^+^ vessels and CD41^+^ megakaryocytes (n = 3 sternum sections from 3 mice). **d,** CD150 and CD71 expression in the indicated HSPC. **e,** FACS plots and colony-forming assays showing that Lin^-^CD117^+^Sca1^-^CD41^-^CD16/32^-^ CD105^+^CD150^+^ Pre CFU-E gated as in Extended Data Fig. 1a are heterogeneous and contain CD71^-^ and CD71^+^ cells with the later fraction containing almost all erythroid colony-forming potential. The data also shows that CD71^+^Ly6C^+^ myeloid progenitors lack erythroid differentiation potential (n = 3 in 3 independent experiments). **f,** FACS plots showing that mature hematopoietic cells do not coexpress CD117 and CD71 or CD117 and CD150. **g-j,** FACS gating strategies for isolation of -and representative images- of erythroid progenitors (g, h) and terminal erythropoiesis (i, j). **k,** Frequencies for the indicated 8 populations in bone marrow by FACS (white) or image (orange) (n = 3 mice for each group). **l-m,** FACS gating strategies for isolation (l) -and representative images (m) - of B lymphopoiesis (the Lin panel contains CD2, CD3e, CD5, CD8, CD11b, Ter119, Ly6G, IgM, and IgD). All data represent individual values with median plot. Statistical differences were calculated using two-tailed unpaired Student’s t-test; *P* values are shown.

**Extended Data Figure 3. Mapping multipotent hematopoiesis and megakaryopoiesis in whole mounted sternum. a,** Experimental workflow for automatic cell segmentation. Fluorescence in the individual channels is merged, converted to 12-bit data, and intensities normalized. The “3D darkspot detection” algorithm detects cells of different sizes. We then watershed each dark centroid to the bright membrane. This is repeated across all z sections until each cell is annotated as an individual object. The generated “inside cell” binary data was exported from Nikon Elements software to Bitplane Imaris software and used to place dots representing each hematopoietic cell. The upper histogram shows sensitivity (= number of correctly segmented cells divided by the number of cells identified manually) and specificity (= number of correctly segmented cells divided by the number of identified cells using the automatic algorithm) of cell segmentation (n = 6 areas in 2 sternum segments from 2 mice). The lower histograms compare the distribution of distances between randomly selected cells segmented manually or through the automatic algorithm. **b,** Histograms showing the distribution of distances from each HSPC to the closest indicated cell (n = 66 ST-HSC, 30 MPP2/MyE, 41 MPP3/MPP4/CMP, 61 MkP, 82 Pre Meg-E/Pre CFU-E in 5 sternum segments from 4 mice). **c.** Distance analyses from each indicated cell to the closest sinusoid, arteriole, or megakaryocyte (n = 42 LT-HSC, 63 ST-HSC, 32 MPP2/MyE, 55 MPP3/MPP4/CMP, 117 MkP, 85 Pre Meg-E in 5 sternum segments from 3 mice). **d,** Map showing the distribution of MkP and megakaryocyte (MK) in a sternum segment. **e,** Experimental design, and histogram showing the percentage of CFP, GFP, RFP, and YFP positive cells in bone marrow megakaryocytes in tamoxifen-treated *Ubc-cre^ERT2^:confetti* mice (each dot represents one sternum segment from 3 confetti mice). **f,** Representative image showing confetti-labeled megakaryocyte progenitors (MkP) and megakaryocytes (MK) in tamoxifen-treated confetti mice. **g,** Observed (Ob) and Random cell (Rd) distances from each confetti-labeled MkP or MK to the closest MK labeled in the same color (n = 23 labeled MkP, 308 labeled MK in 3 sternum segments from 3 tamoxifen-treated confetti mice). Statistical differences were calculated using two-tailed unpaired Student’s *t*-tests; *P* values are shown.

**Extended Data Figure 4. Mapping erythropoiesis, B Lymphopoiesis and myelopoiesis in whole mounted sternum. a**, Distance analyses from each erythroid progenitor to the closest arteriole (n = 111 Pre Meg-E, 18 Pre CFU-E, and 523 CFU-E in 3 sternum segments from 3 mice). **b,** Distance analyses from CFU-E to the closest indicated cell (n = 318 CFU-E in 3 sternum segments from 3 mice). **c,** Distance analyses from erythroblasts (CD117^-^CD71^+^Ly6C^-^, containing early and late erythroblasts, n = 4461 in 3 sternum segments from 3 mice) to the closest indicated erythroid progenitor. **d,** Representative images and distance analyses of confetti-labeled CFU-E to the closest CFU-E labeled in the same color (n = 38 confetti-labeled CFU-E in 5 sternum segments from 3 tamoxifen-treated confetti mice). **e,** Representative image showing confetti-labeled erythroblasts (CD117^-^CD71^+^Ly6C^-^, containing early and late erythroblasts) in tamoxifen-treated confetti mice. Due to a lack of available fluorescence channels for analyses, Ly6C and GFP were combined in a single dump channel. The histogram shows distance analyses of each confetti-labeled erythroblast to the closest erythroblast labeled in the same color (CD117^-^CD71^+^Ly6C^-^, containing early and late erythroblasts, n = 267 labeled erythroblasts in 4 sternum segments from tamoxifen-treated confetti mice). **f,** FACS gating strategy for simultaneous detection of CD117^+^CD127^+^Lin-mix^-^CLP and CD117^+^ESAM^+^ HSPC (the Lin-mix panel contains B220, CD2, CD3e, CD5, CD8, CD11b, Ter119, Ly6G, CD41, Ly6C). **g,** Representative image and distance analyses showing that CLP map near arterioles and away from sinusoids (n = 36 CLP in 3 sternum segments from 3 mice). **h,** Representative image and distance analyses of confetti-labeled CLP or B cell precursors (B220^+^Lin^-^ cells, the Lin panel contains CD2, CD3e, CD5, CD8, CD11b, Ter119, Ly6G, IgM, and IgD, containing all Pre-pro B, Pro B, and Pre B cells) to the closest B cell precursors labeled in the same color (n = 11 confetti-labeled CLP, 23 confetti-labelled B cell precursors in 3 sternum segments from 3 tamoxifen-treated confetti mice). **i,** Map showing monocyte dendritic progenitor cell (MDP) and granulocyte progenitor (GP) localization to sinusoids. **j, k,** Representative images showing a DC/Ly6C^lo^ monocytes production line around an MDP (j), and a neutrophil production line centered around a GP (k). In all maps dots are three times the average size of the relevant cell. **l,** Map showing the distribution of Pre CFU-E / CFU-E –labeling erythroid production lines, Pre and immature neutrophils –labeling neutrophil production lines, and Ly6C^lo^ Monocytes (Ly6C^lo^ Mo, labeling cDC/ Ly6C^lo^ monocyte production lines) in the sternum. Map dots are two times the average size of the relevant cells. **m,** Distance analyses from each CFU-E to the closest indicated cells (PN + IN: Pre and immature neutrophils; n = 297 CFU-E in 3 sternum segments from 3 mice). n, Quantification of each type of production line per sternum segment (n = 4 sternum segments from 4 mice). Statistical differences were calculated using two-tailed unpaired Student’s t-tests; *P* values are shown.

**Extended Data Figure 5. Quantification of bone marrow cell number changes in mice during acute stress. a, b**, Number of the indicated cells per femur (detected by FACS. Bone marrow HSPC populations are identified as in Extended Date Figure 1.a; granulopoiesis and Mo- and DC-poiesis populations are identified as described in reference^1^; erythropoiesis populations are identified as described in Extended Date Figure 2 g and i; lymphopoiesis populations are identified as described as in Extended Date Figure 2 l) at the indicated time points after phlebotomy (a) or *Listeria monocytogenes infection* (b). Each dot corresponds to one mouse in 3-6 experiments (n = 4-12 mice for each time point). All data represent individual values with median plot. Statistical differences were calculated using two-tailed unpaired Student’s *t*-tests; *P* values are shown.

**Extended Data Figure 6. The basic anatomy of hematopoiesis is largely maintained during acute stress. a,** Scheme showing the overall experiment design (Top panels) and quantification (bottom panels) of the indicated populations (normalized to day 0) in response to *L. monocytogenes* infection (n = 4 mice for each time point). All data represent mean ± s.e.m. **b-d,** Vascular organization in control (con), phlebotomized (phl), and *L. monocytogenes-*infected (inf) mice. Map (b); vessel numbers (n = 3, control and phlebotomy, or n = 4, infection, sternum segments from 3 mice), sinusoid length and diameter (c, n = 50 randomly selected sinusoids from 3 sternum segments from 3 mice); and arteriole and megakaryocyte numbers (d, n = 4, control, phlebotomy, or infection sternum segments from 4 mice). **e,** Distance analyses from each HSPC to the closest indicated cells at the indicated time points after challenge. (n = 41 LT-HSC, 66 ST-HSC, 30 MPP2/MyE, 41 MPP3/MPP4/CMP, 61 MkP, and 82 Pre Meg-E in 4 sternum segments from 4 mice in steady-state for control; n = 21 LT-HSC, 35 ST-HSC, 16 MPP2/MyE, 30 MPP3/MPP4/CMP, 38 MkP, and 73 Pre Meg-E in 3 sternum segments from 3 mice two days after phlebotomy; n = 15 LT-HSC, 19 ST-HSC, 56 MPP2/MyE, 39 MPP3/MPP4/CMP, 17 MkP, and 57 Pre Meg-E in 3 sternum segments from 3 mice six days after infection). **f,** Representative image showing a loose MPP2/MyE cluster**. g-i,** Maps (g) and distance analyses showing CLP localization to sinusoid (h, steady-state (d0) n = 36, and six days after infection (d6) n = 43 CLP in 3 sternum segments from 3 mice) and CFU-E localization to sinusoids (i, steady-state (d0) n = 315, and six days after infection (d6) n = 314 CFU-E in 3 sternum segments from 3 mice). **j,** Maps showing MDP and GP distribution in the sternum after stress. **k-n**, Representative images (k, l) and quantification (m, n) showing transient GP and MDP detachment from sinusoids after infection. (n = 62, 16, 18 MDP; and n = 114, 98, 49 GP in 3-5 sternum segments from 3 mice in steady-state (d0), six days (d6), and twenty days (d20) after infection). **o,** Quantification of each type of production line per sternum after infection (n = 3-5 sternum segments from 3-4 mice for each time point). All data represent mean ± s.e.m. Dots in maps are three times the size of the relevant cell except for CLP dots which are five times the actual size. Statistical differences were calculated using two-tailed unpaired Student’s *t*-tests; *P* values are shown.

**Extended Data Figure 7. Production lines orchestrate hematopoietic plasticity to stress. a, b,** Representative images (a) and number of cells per B cell production line (b) from control or *L. monocytogenes*-infected mice after infection (n = 12 randomly selected production lines in 3 sternum segments from 3 mice for each time point). **c,d,** Map and representative images showing the changes in erythroid production lines after infection. Map dots are three times the average size of the relevant cell. **e**, Number of CFU-E (n = 65, 45, and 60 production lines for days 0, 6, 20, three sternum segments from 3 mice per time point) and indicated erythroid cells (n = 5 randomly selected production lines for each time point, in 3 sternum segments from 3 mice) from control or *L. monocytogenes*-infected mice. **f-h,** Maps (f), representative images (g) and number of cells (h) in DC/Ly6C^lo^ monocyte production lines from control or *L. monocytogenes*-infected mice (n = 49, 35, and 29 randomly selected production lines for days 0, 6, 20, in 3-5 sternum segments from 3 mice per time point). **i, j,** as g-h but for granulocyte production lines (n = 96, 88, and 50 randomly selected production lines in 3-5 sternum segments from 3-4 mice per time point). Dots in maps are three times the size of the relevant cell. Statistical differences were calculated using two-tailed unpaired Student’s *t*-tests; *P* values are shown.

**Extended Data Figure 8. Quantification of bone marrow cell number changes in aged mice. a,** Number of BM cells or the indicated populations per femur (detected by FACS. Bone marrow HSPC populations are identified as in Extended Date Figure 1.a; granulopoiesis and Mo- and DC-poiesis populations are identified as described in reference^1^; erythropoiesis populations are identified as described in Extended Date Figure 2.g and i; lymphopoiesis populations are identified as described as in Extended Date Figure 2.l) in 2-month-old (n = 7) and 20-month-old (n = 4 - 10) mice). Each dot corresponds to one mouse in 3 independent experiments. All data represent individual values with median plot. Statistical differences were calculated using two-tailed unpaired Student’s *t*-tests; *P* values are shown.

**Extended Data Figure 9. The basic anatomy of hematopoiesis is maintained in aged mice. a,** Histograms showing CD41 and CD42d expression in the indicated HSPC in 2- and 20-month old mice. **b,** FACS plots showing the gating strategy to identify the indicated HSPC based on CD42d and CD41 in 20-month-old mice. **c, d,** Representative images (c) and quantification of cell diameters (d) demonstrating that ESAM^+^CD117^+^CD48^-^CD150^+^CD42^-^CD41^+^ myeloid-biased LT-HSC, ESAM^+^CD117^+^CD48^-^CD150^+^CD42^-^CD41^-^ lymphoid-biased LT-HSC and ESAM^+^CD117^+^CD150^+^CD42^+^CD41^+^ MkP can be distinguished based on cell size (n = 30 randomly selected cells from each type in 3 sternum segments from three 20-month-old mice). **e,** Sensitivity (= Number of ESAM^+^CD117^+^CD48^-^CD150^+^CD41^-^ LT-HSC, ESAM^+^CD117^+^CD48^-^CD150^+^CD41^+^ LT-HSC, ESAM^+^CD117^+^CD150^+^CD41^+^ MkP correctly identified based on CD150 and CD41 expression and cell size, divided by the number of cells identified when counterstained with CD42 (CD41^-^CD42^-^ LT-HSC, CD41^+^CD42^-^ LT HSC, and CD41^+^CD42^+^ MkP) and specificity (= Number of ESAM^+^CD117^+^CD48^-^CD150^+^CD41^-^ LT-HSC, ESAM^+^CD117^+^CD48^-^CD150^+^CD41^+^ LT-HSC, ESAM^+^CD117^+^CD150^+^CD41^+^ MkP correctly identified based on CD150 and CD41 expression and cell size, divided by the number of cells identified based on CD150 and CD41 expression and cell size) for distinguishing myeloid and lymphoid biased LT-HSC and MkP based on CD41 and CD150 expression and cell size (compared with CD42d based identification, 47 CD41^+^ LT-HSC, 55 CD41^-^ LT- HSC, and 70 MkP in n = 3 sternum segments from two 20-month-old mice were analyzed). **f-h,** Age- dependent changes in the vascular microenvironment. Maps (f), number of sinusoids, arterioles, and megakaryocytes (g, n = 3 sternum segments from three mice for vessel quantification and n = 4 sternum segments from four 2- and 20-month-old mice for megakaryocyte quantification); and sinusoid length, diameter, and branching points (h, n = 50 sinusoids randomly selected in 2 sternum segments from two 2- and 20-month-old mice). **i,** Distance analyses from each HSPC to the closest indicated cells in 20-month-old mice. (n = 191 CD41^-^ LT-HSC, 109 CD41^+^ LT-HSC, 236 ST-HSC, 39 MPP2/MyE, 72 MPP3/MPP4/CMP, 133 MkP and 57 Pre Meg-E in 5 sternum segments from three 20- month-old mice). **j**, Distance analyses from each HSPC to the closest sinusoid and arteriole. (n = 315 ST-HSC, 46 MPP2/MyE, 102 MPP3/MPP4/CMP, 131 MkP and 79 Pre Meg-E in 5 sternum segments from three 20-month-old mice). **k**, Maps showing committed progenitor distribution in the aged marrow. **l**, Distance analyses from each committed progenitor to the closest sinusoid or arteriole (n = 558 CFU-E, 115 GP, 33 MDP in 3 sternum segments from three 20-month-old mice). **m**, Distance analyses showing a lack of CLP localization to arterioles or sinusoids in 20-month-old mice (n=38 CLP in three sternum segments from three 2- and 20-month-old mice) in aged mice. Dots in maps are 3-times (CFU-E, GP, and MDP), or 5-times (CLP) the size of the relevant cell. **n,** Representative images showing an LT-HSC cluster in the aged sternum. Statistical differences were calculated using two-tailed unpaired Student’s *t*-tests; *P* values are shown.

**Extended Data Figure 10. Aging caused extensive cell and lineage-specific microanatomical adaptations in each type of production line. a,** Representative images showing stage-specific delayed B cell differentiation (the Lin panel contains CD2, CD3e, CD5, CD8, CD11b, Ter119, Ly6G, IgM, IgD). **b-d,** Maps (b) and representative images showing delayed differentiation in granulocyte production lines (c) and increased dendritic cell and reduced monocyte output (d) in dendritic cell (DC)/Ly6C^lo^ monocytes (Ly6C^lo^ Mo) production lines in aged mice. Dots in maps are 3-times (GP), or 2-times (PN) the size of the relevant cell.

## Methods

### Mice

All mouse experiments were approved by the Institutional Animal Care Committee of Cincinnati Children’s Hospital Medical Center. The following mouse strains were used: C57BL/6J*-Ptprc^b^* (CD45.2), B6.SJL-*Ptprc^a^Pepc^b^*/BoyJ (CD45.1), B6.Cg*-Ndor1^Tg(UBC-cre/ERT2)1Ejb^/1J* (Ubc-cre^ERT2^) and B6.129P2-*Gt(ROSA)26Sor^tm1(CAG-Brainbow2.1)Cle^* (R26R-Confetti). R 26R-Confetti mice were crossed with Ubc-cre^ERT2^ mice to generate *Ubc-cre^ERT2^:Confetti* mice. All mice were maintained on a C57BL/6J background. Eight to twelve (2-month-old) and eighty to a hundred weeks (20-month-old) male and female mice were used. All mice were bred and aged in our vivarium or purchased from the Jackson Laboratory. Mice were maintained at the vivarium at Cincinnati Children’s Hospital Medical Center under a 14-hours light:10-hours darkness schedule, 30–70% humidity, 22.2 ± 1.1 °C, and specific-pathogen-free conditions.

### Tamoxifen treatment

*Ubc-cre^ERT2^:Confetti* mice were treated with two pulses of tamoxifen in the diet (400 mg of Tamoxifen Citrate/Kg of rodent diet, Envigo). Each pulse was two weeks long and pulses were two weeks apart. Since committed hematopoietic progenitors do not persist in vivo for longer than two weeks we chased the mice for eight weeks to ensure that all confetti-labeled immature and mature hematopoietic cells originated from upstream progenitors.

### Listeria monocytogenes infection

The wild-type virulent *L. monocytogenes* strain 10403s was back-diluted from overnight culture for 2 hours to early log phase growth (OD_600_ 0.1) in BD Difco™ brain-heart infusion media (Thermo Fisher Scientific, catalogue number 237500) at 37°C, then washed and diluted in 200 μl sterile saline and injected via the lateral tail vein to mice (1 x 10^4^ CFUs per mouse). Mice were sacrificed for bone marrow analyses on day 6 and 20 after infection.

### Phlebotomy mice model

To induce erythropoietic stress by blood loss, isoflurane-anesthetized mice were phlebotomized (15-20 μl blood/gram of body weight from the retro-orbital venous sinus of mice) with a calibrated heparinized capillary tube. Mice were euthanized for bone marrow analyses on day 2, 8, and 20 after phlebotomy.

### Cell preparation for flow cytometry and cell sorting

Mice anesthetized with isoflurane followed by cervical dislocation. Bone marrow cells were flushed out of the femurs with a 21-gauge needle in 1 ml of ice-cold PEB buffer (2 mM EDTA and 0.5% bovine serum albumin in PBS). Peripheral blood was collected from the retro-orbital venous sinus of mice, followed by red blood cell lyses with 1 ml of lysis buffer (150 mM NH4Cl, 10 mM NaCO3, and 0.1 mM EDTA). Cells were centrifuged for 5 minutes at 1100 rpm under 4 °C, resuspended in ice-cold PEB, and used in subsequent assays. For FACS cell sorting, the bone marrow cells were stained with a cocktail of biotinylated lineage antibodies for 30 minutes, washed twice, and stained with streptavidin-conjugated magnetic beads (BD Bioscience, catalogue number 557812). Magnetic cell depletion was performed according to the manufacturer’s protocol. CountBright™ Absolute Counting Beads (Thermo Fisher Scientific, catalogue number C36950) were used to count bone marrow and blood cell numbers in a BD LSRFortessa™ Flow Cytometer (BD Bioscience).

### FACS analyses and LEGENDScreen

Cells were stained in the dark for 30 minutes in ice-cold PEB buffer containing antibodies, washed thrice with ice-cold PEB, and analyzed in an LSRFortessa™ Flow Cytometer (BD Biosciences), LSR II Flow Cytometer (BD Biosciences), or FACS-purified in a FACSAria™ II Cell Sorter (BD Biosciences) or an SH800S Cell Sorter (Sony Biotechnology). Dead cells and doublets were excluded on the basis of FSC and SSC distribution and DAPI exclusion (Sigma-Aldrich, catalogue number D9542) exclusion. Antibodies used were: B220 (clone RA3-6B2), CD2 (clone RM2-5) CD3e (clone 145-2C11), CD4 (clone RM4-5), CD5 (clone 53-7.3), CD8 (clone 53-6.7), CD11b (clone M1/70), CD11c (clone N418), CD16/32 (clone 93), CD24 (clone 30-F1), CD31 (clone A20), CD41 (clone MWReg30), CD42d (clone 1C2), CD43 (clone S11), CD45 (clone 30-F11), CD45.1 (clone A20), CD45.2 (clone 104), CD48 (clone HM48-1), CD71 (clone RI7217), CD105 (clone MJ7/18), CD115 (clone AFS98), CD127 (clone A7R34), CD135 (clone A2F10), CD144 (clone BV13), CD150 (clone TC15-12F12.2), ESAM (clone 1G8), Gr1 (clone RB6-8C5), IgD (clone 11-26c.2a), IgM (clone RMM-1), Ly-6C (clone HK1.4), Ly-6G (clone 1A8), Sca-1 (clone D7), Ter119 (clone TER-119), MHC II (clone M5/114.15.2), from BioLegend; CD34 (clone RAM34) and CD117 (clone 2B8), from BioLegend or Thermo Fisher Scientific; CD71 (clone C2) from BD Bioscience. For immunophenotyping experiments, LEGENDScreen™ Mouse PE Kit (BioLegend, catalogue number 700005) was used according to the manufacturer’s instructions. Briefly, fresh bone marrow cells were stained with a cocktail of biotinylated lineage antibodies for 30 min followed by a stain with streptavidin. Cells were washed twice and resuspended at a concentration of 1 × 10^7^ cells per ml PEB buffer containing antibodies for hematopoietic stem and progenitor cells identification. Equal amount of cells were transferred into each well of the LEGENDScreen 96-well plates. Plates were incubated for 45 minutes on ice in the dark. Cells were then washed twice and resuspended in PEB buffer and kept on ice until acquisition on a BD LSRFortessa™ Flow Cytometer (BD Bioscience). FACS Data was analyzed with FlowJo software (Tree Star). Gating strategies for all analyses are shown in the main or Extended Data Figures.

### Colony-forming unit (CFU) assay

FACS-purified cells were suspended in IMDM + 2% FBS, added into the methylcellulose culture medium (StemCell Technologies, MethoCult™ M3334, M3434, M3436, and M3534,), mixed thoroughly via vortex, plated in duplicate 35 mm culture dishes (Greiner Bio-One, Catalog #627160), and incubated at 37°C with 5% CO2 in air and ≥ 95% humidity, for 7-10 days. Colonies were identified and counted based on cluster size and cell morphology using Nikon Eclipse Ti inverted microscope (Nikon Instruments Inc., NY, USA) equipped with 4x, 10x, and 40x objectives.

### Competitive LT- and ST-HSC transplantation

Adult CD45.1^+^ recipient mice were lethally irradiated (700 rads plus 475 rads, 3 hours apart). The indicated number of FACS purified CD45.2^+^ HSCs was mixed with 10^6^ CD45.1^+^ competitor mouse bone marrow cells and transplanted by retro-orbital venous sinus injection within 6 hours after the second irradiation. Peripheral blood chimerism was determined by FACS analyses at week 4, 8, 12, and 16 post-transplant.

### Transplant of ESAM^+^ and ESAM^-^ progenitor subsets in sublethally irradiated recipients

Adult CD45.1^+^ recipient mice were sublethally conditioned with a single dose of 700 rads. The indicated number of FACS-purified ESAM^+^ or ESAM^-^ HSPC were transplanted via retro-orbital venous sinus injection within 6 hours after irradiation. Peripheral blood chimerism was determined by FACS analyses on day 10, 20, 30, and 40 post-transplant.

### Whole-mount immunostaining

In experiments requiring visualization of blood vessels, mice were intravenously injected with 10 μg of Alexa Fluor 647 anti-mouse CD144 antibody (BV13, BioLegend) and euthanized 10 minutes after injection. In experiments requiring visualization of CLP, mice were intravenously injected with 2 μg of Alexa Fluor 647 anti-mouse CD127 antibody (A7R34, BioLegend) and euthanized 5 minutes after injection. Whole-mount sternum immunostaining has been described before^30^. Briefly, the sterna were dissected and cleaned of soft and connective tissue, followed by sectioning along the sagittal or coronal plane to expose the bone marrow under a dissecting microscope (Nikon SMZ1500 Stereomicroscope). Each half of the sternum was fixed in 4% PFA (Electron Microscopy Sciences, catalogue number 15710) in DPBS (Thermo Fisher Scientific, catalogue number 14190144) for 3 hours on ice. Each fragment was further washed with DPBS after fixation and blocked with 10% goat serum (Sigma-Aldrich, catalogue number G9023) for 1 hour, followed by staining with 100 µl staining buffer (2% goat serum in DPBS and the indicated antibodies) on ice.

### Confocal imaging

Confocal imaging was performed in a Nikon A1R GaAsP Inverted Confocal Microscope, Nikon A1R LUN-V Inverted Confocal Microscope, or Nikon AXR Inverted Confocal Microscope. Specifications for the Nikon A1R GaAsP Inverted Confocal Microscope: high power 405 nm, 442 nm, 488 nm, 561 nm, 640 nm, and 730 nm solid-state diode lasers. Specifications for the A1R LUN-V Inverted Confocal Microscope: high power 405 nm, 445 nm, 488 nm, 514 nm, 561 nm, and 647 nm solid-state diode lasers. Specifications for the AXR Inverted Confocal Microscope: high power 405 nm, 445 nm, 488 nm, 514 nm, 561 nm, 594 nm, 640 nm, and 730 nm solid-state diode lasers. All microscopes were equipped with a fully encoded scanning XY motorized stage, Piezo-Z nosepiece for high-speed Z-stack acquisition, resonant and galvanometric scanners, one high-quantum efficiency, low-noise Hamamatsu photomultiplier tube, and three high-quantum efficiency gallium arsenide phosphide photomultiplier tubes (GaAsP-PMTs) for overall 400-820nm detection. An LWD Lambda S 20XC water-immersion objective (Nikon, MRD77200) was used and images were taken using the resonant scanner with 8x line averaging, 1024 x 1024 pixels resolution, and 2 μm Z-step. For high power images we used a LWD Lambda S 40XC water-immersion objective (Nikon, MRD77410) with a resonant scanner and 8X line averaging, 1024 x 1024 pixels resolution, 0.5 μm Z-step.

### Image and distance analyses

Original images (.ND2 format file) were denoised by a built-in artificial intelligence algorithm (Denoise.AI) and stitched together using the NIS-Elements software (Nikon, version 5.20.02 and 5.30.03). The denoised and stitched ND2 files were converted to Imaris (.IMS) files using Imaris software (Bitplane, version 9.5 to 9.9). Because not all antibodies penetrate to the same depth within the tissue, we only examine the first 35 µm of the sternum image, which we have previously shown are uniformly stained through the tissue^1^. Cells of interest were labeled with dots with the Imaris Spots function in manual mode and dots X, Y, and Z coordinates were automatically computed. Sinusoids, arterioles, and megakaryocytes were segmented based on channels of CD144, CD41, ESAM, and Ly6C using the Imaris Surface function. The diameters of each type of cell were measured manually in 3D view in Imaris software and were as follows: CD41^-^ LT-HSC, 8.67 ± 1.23 μm; CD41^+^ LT-HSC, 8.94 ± 0.91 μm; ST-HSC, 8.68 ± 1.10 μm; MPP2, 7.98 ± 1.05 μm; MPP3, 8.48 ± 1.32 μm; MkP, 14.45 ± 3.88 μm; Pre Meg-E, 9.49 ± 1.34 μm; Pre CFU-E, 13.92 ± 1.70 μm; CFU-E, 12.67 ± 1.88 μm; Early Erythroblast, 8.86 ± 1.61 μm; Late Erythroblasts, 7.92 ± 1.36 μm; Reticulocytes, 5.17 ± 0.76 μm; RBC, 4.38 ± 0.60 μm; CLPs, 7.40 ± 0.97 μm; Pre-pro B, 8.9 ± 0.61 μm; Pro B, 7.71 ± 1.23 μm; Pre B, 6.10 ± 0.61 μm; MDP, 12.13 ± 1.19 μm; GP, 11.70 ± 0.99 μm; PN, 10.21 ± 1.08 μm; Ly6C^lo^ Mo, 9.30 ± 1.17 μm; cDC, 12.33 ± 2.69 μm. The distance from each cell to the closest vascular structures and megakaryocytes was obtained with the Imaris Distance Transform Matlab Xtension and then subtracted the mean radius for each cell type. The distance between cells was calculated using Matlab software (MathWorks, version 2018a) with the coordinates exported from Imaris and then subtracted the mean radius for each cell. All software were installed in HP Z4 windows 10 x 64 workstations equipped with Dual Intel Xeon Processor W-2145, 192GB ECC-RAM, and an Nvidia Quadro RTX 5000 16GB GDDR6 Graphics Card.

### Confetti imaging

For our imaging experiments we have use six fluorescent channels (405 nm, 445 nm, 488 nm, 514 nm, 561 nm, and 647 nm). In the confetti model Cre recombination leads to expression of GFP (488 nm), YFP (514 nm), RFP (561 nm), and CFP (445 nm) thus occupying four out of six channels used for imaging. To overcome this limitation and analyze spatial relationships between confetti labeled cells we routinely used a dump channel with Alexa 488 or FITC-labeled antibodies (same fluorescence as GFP). We discarded cells showing green fluorescent from analyses and compared YFP, RFP and CFP labeled cells of interest. To analyze the clonal relationships between MkP and megakaryocytes, we used CD41-FITC that spectrally overlap with GFP. We discarded CD41^+^GFP^+^ cells from analyses (Extended Data Fig. 3f). To analyze the clonal relationships between CFU-E and erythroblasts, we used Ly6C-Alexa 488, and discarded Ly6C^+^GFP^+^ cells from analyses (Extended Data Fig. 4d, e). To analyze the clonal relationships between CLP and B precursors, we used Lin-Alexa 488 (the Lin panel contains CD2, CD3e, CD5, CD8, CD11b, Ter119, Ly6G, IgM, IgD), and discarded Lin^+^ GFP^+^ cells from analyses (Extended Data Fig. 4h).

### Random simulations

Sternal fragments were stained with anti-CD45 and anti-Ter119 antibodies to detect all hematopoietic cells, with anti-CD144, anti-ESAM, anti-CD41 and anti-Ly6C antibodies to detect sinusoids, arterioles, and megakaryocytes. 3D binary segmentation tools in NIS Elements software were used to automatically annotate CD45^+^ or Ter119^+^ cells. Briefly, high-resolution images (0.31 um/pixel XY, 0.6 um/pixel Z) acquired with a 40x Water Immersion objective (NA 1.15) were deconvolved, and CD45 and Ter119 fluorescent membrane channels were added into a single channel with the floating-point math, converted into 12-bit data, and pre-processed to normalize intensities in-depth and min/max intensities. The “3D darkspot detection” algorithm allows the detection of cells of different sizes. This segmentation algorithm considers the distribution of intensities in X, Y, and Z. 3D region watershed dark centroid to bright membrane. This will account for non-spherical cells and include all dark space inside the cell membrane stain. The generated “inside cell” binary data was exported to the Imaris software and used to place dots representing each hematopoietic cell (48964 to 81248 cells) in each 35 µm optical slice Z-stack of each sternum fragment. We then used Research Randomizer^30^ to randomly select dots representing each type of hematopoietic cell at the same frequencies found in vivo through the bone marrow cavity and measured the distances between these random cells or with vessels as above. Each random simulation was repeated 100-200 times.

For generating random distributions of cells in experiments using confetti mice we first obtained the coordinates and confetti color for each type of cell in each section analyzed. Then we used Research Randomizer ^31^ to randomize the confetti label while maintaining the spatial coordinates of each cell. We then measured the distances between these cells with randomized colors. Each random simulation was repeated 100-200 times.

### Defining the anatomy of each lineage-specific production line

Our analyses comparing the observed and random distances from each type of late erythroid cells to CFU-E (Fig. 2h) showed that these cells were much closer to CFU-E than random cells. The mean of random cells to CFU-E was 46.18 µm for early erythroblasts, 48.33 µm for late erythroblasts, 48.98 µm for reticulocytes, and 49.98 µm for erythrocytes (Fig. 2h). We thus defined production lines as the region within 50 µm of a sinusoids CFU-E string.

Our analyses comparing the observed and random distances from each type of B precursors (including Pre-proB, Pro B and Pre B) to CLP (Fig. 2n) showed that these cells were much closer to CLP than random cells. The mean of random cells to CLP was 135.5 µm for Pre-pro B, 136.1 µm for Pro B, and 161.9 µm for Pre B (Fig. 2n). We thus defined B cell production lines as the region within 150 µm of a CLP. The granulocyte production lines, and dendritic cell (DC) / Ly6C^lo^ monocyte (Ly6C^lo^ Mo) production lines were defined as we described previously^1^.

### Quantify vessel length, diameter and branching

Bone marrow vessels were detected based on ESAM and Ly6C expression (sinusoids ESAM+Ly6C^-^, arterioles ESAM^+^Ly6C^+^). We defined a branch as the point where two or more lumens connect. A vessel is a vascular structure -with a continuous lumen-between two branching points. Vessel length and diameter were measured manually using the “measurement” tool in Imaris. Diameter reported was the largest value for the whole vessel.

### Statistics

All statistical analyses were performed using Prism 9 (GraphPad Software). For graphs quantifying cells in different mice, we indicate the mean, and each dot corresponds to one mouse. For graphs showing distances between cells and structures each shape corresponds to one cell and different shapes correspond to cells in different mice. The blue bar indicates the median. For graphs showing peripheral blood chimerism determination by FACS analyses after multipotent hematopoietic progenitors’ transplant, ESAM^-^ group compared with ESAM^+^ group at every indicated time point.

Statistical analysis was performed using two-way ANOVA followed by Sidak’s multiple comparisons test. For graphs of stress model data analyses, statistical differences between stress model and steady-state for every indicated time point were calculated using multiple two-tailed unpaired Student’s *t*-tests. All two group comparisons were calculated using two-tailed unpaired Student’s *t*-tests. \**p* < 0.05; \*\**p* < 0.01; \*\*\**p* < 0.001; \*\*\*\**p* < 0.0001; NS, not significant.

### Data reporting

No statistical methods were used to predetermine sample size. All mice were included in the analyses. Mice were randomly allocated to the different groups based on the cage, genotype, and litter size. For all experiments, we aimed to have the same number of mice in the control and experimental groups. Investigators were not blinded to allocation during experiments and outcome assessment.

## Acknowledgements

We thank the Confocal Imaging Core, the Research Flow Cytometry Core and the Veterinary Services at the University of Michigan and Cincinnati Children’s Medical Center for experimental and technical assistance. This work was supported by the National Heart Lung and Blood Institute. We are deeply grateful to Dr. Andres Hidalgo for critical reading of this manuscript.

Daniel Lucas is supported by R01 HL136529, R01 HL153229, R01 HL158616, R01HL160614 and U54 DK126108. Jizhou Zhang is supported by an Evans MDS Foundation Young Investigator Award. Marie-Dominique Filippi is supported by R01 HL151654. H. Leighton Grimes is supported by R01 HL122661. Sing Sing Way is supported by the NIH through award DP1AI131080, the HHMI Faculty Scholar’s Program (grant #55108587), Burroughs Wellcome Fund, and the March of Dimes Foundation Ohio Collaborative. Data was generated using an SH800 cell sorter funded by NIH grant S10OD023410.

## Author contributions

D.L. conceptualized and managed the study. D.L., J.Z., Q.W., J.M.K., and M.D.F designed experiments. Q.W., and J.Z., performed most imaging experiments and analyses. C.B.J., A.S., B.W., M.M., B.S., and H.L.G., maintained mice and collected data. B.E.S., T.Z.S. and S.S.W. performed infection experiments. D.L., Q.W., and J.Z. assembled the figures and wrote the manuscript, with editorial input from all authors.

