## Supplemental Files for "Resilient anatomy and local microplasticity of naïve and stress hematopoiesis"

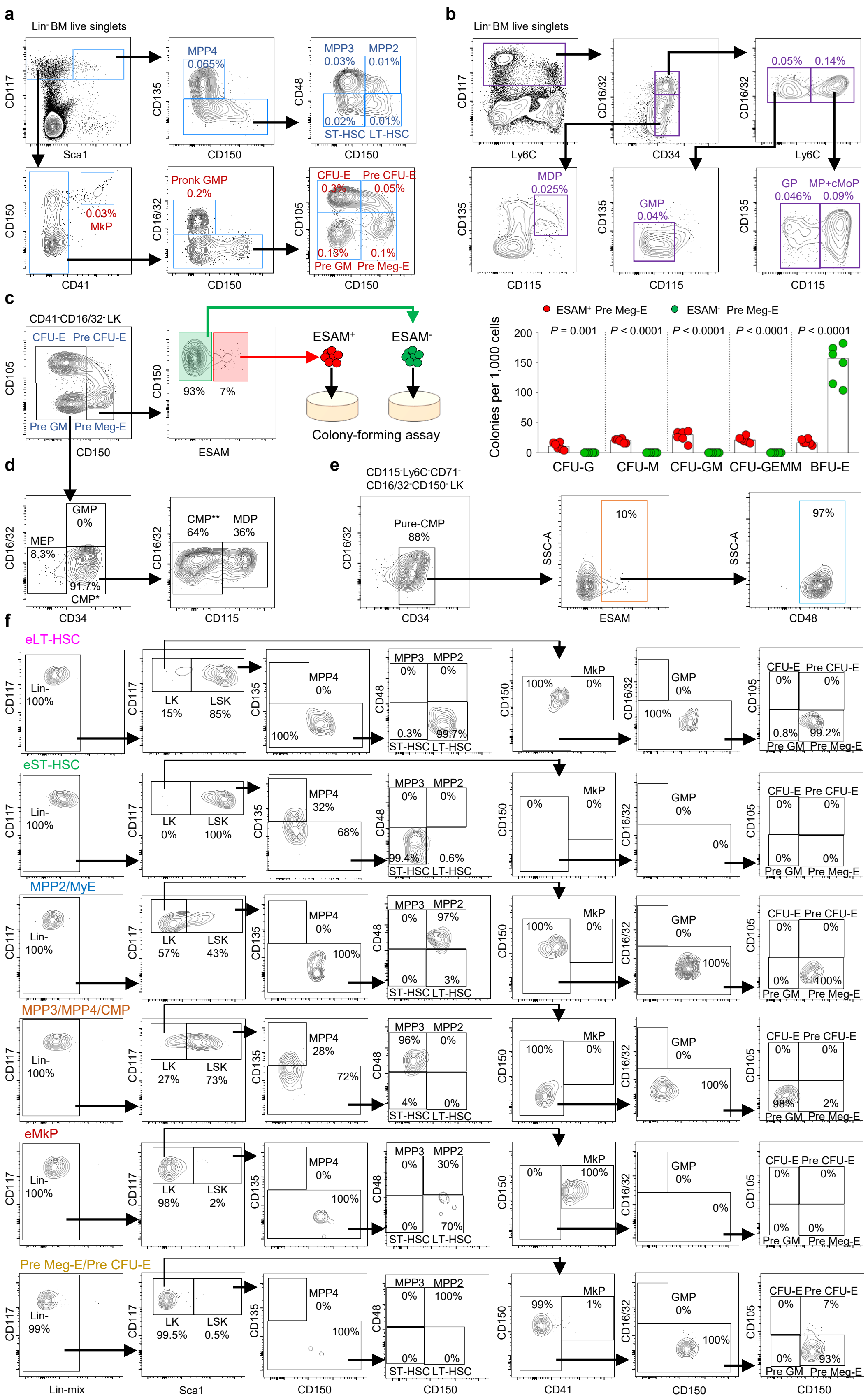


Extended Data Figure 1

**Extended Data Figure 1. Resolving functional heterogeneity of HSPC subsets. a, b,** FACS plots showing the gating strategy to identify 14 indicated hematopoietic progenitors (LT-HSC, ST-HSC, MPP2, MPP3 and MPP4 are identified as described by Pietras et al.[17]; MkP, Pre Meg-E, Pre CFU-E, CFU-E and Pre GM are identified as described by Pronk et al.[14]; GMP, GP, MDP and MP+cMoP are identified as described by Yanez, A. et al.[15] ) that were used for immunophenotyping screen in Fig.1. **c,** FACS plots and colony-forming assays showing that $Lin^-CD117^+Sca1^-CD41^-CD16/32^-CD105^-CD150^+$ Pre Meg-E from (a) are heterogeneous and contain $ESAM^+$ cells with myeloid and erythroid colony-forming activity ($ESAM^+$ MyE) and bona fide Pre Meg-E ($ESAM^-$) (n = 6 from 3 independent experiments). All data represent individual values with median plot. Statistical differences were calculated using multiple two-tailed unpaired Student's t-test; *P* values are shown. **d,** FACS plots showing that the $Lin^-CD117^+Sca1^-CD41^-CD16/32^-CD105^-CD150^-$ Pre GM from (a) are heterogeneous and contain both common myeloid progenitors (CMP) and monocyte dendritic cell progenitors (MDP). **e,** FACS plots showing the gating strategy to identify pure CMP and ESAM and CD48 expression in these cells. **f,** FACS plots showing the relative purity of the HSPC identified using the strategy shown in Fig.1e. (eLT-HSC = $ESAM^+$ LT-HSC; eST-HSC = $ESAM^+$ ST-HSC; eMkP = $ESAM^+$ MkP).

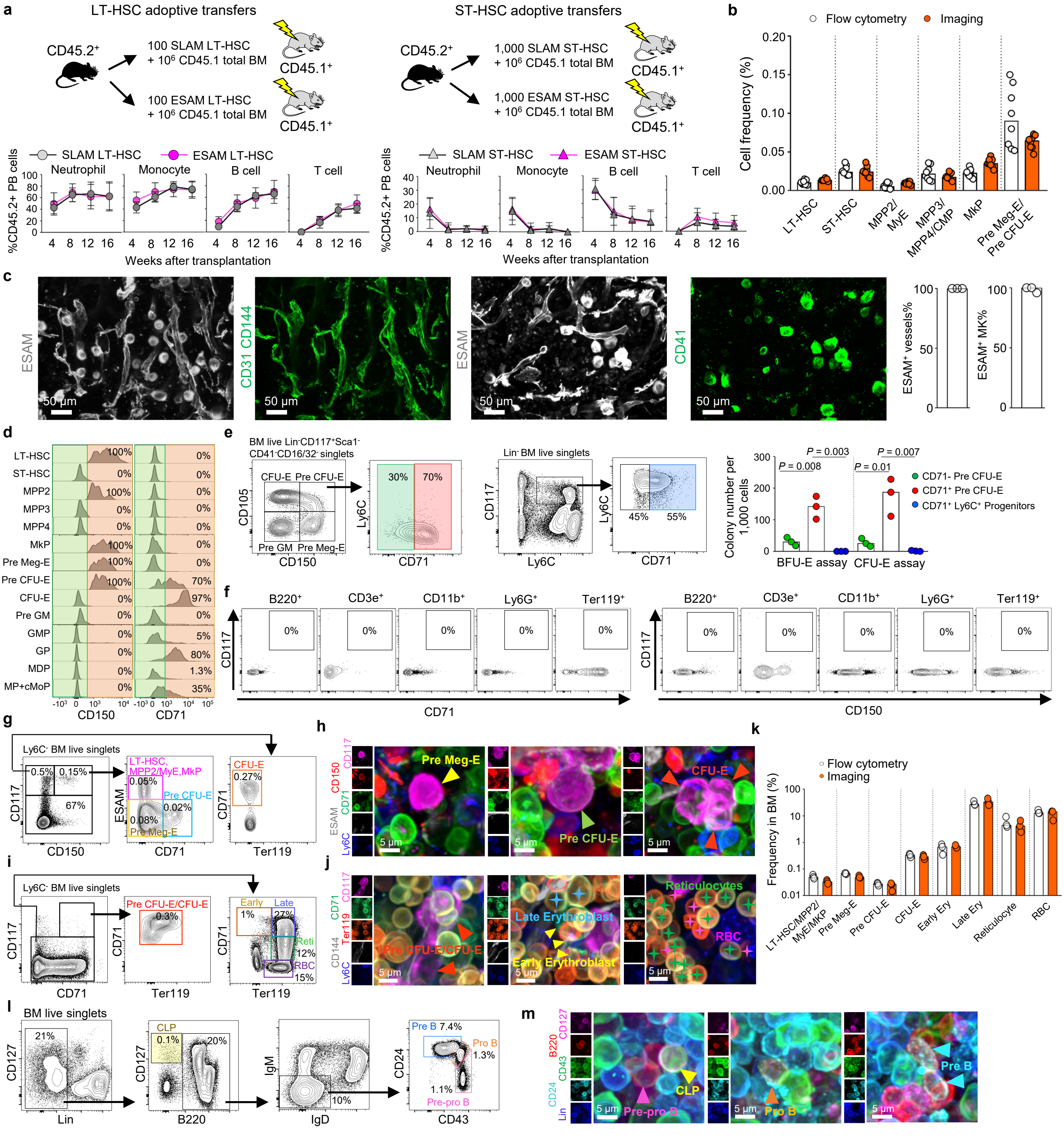


**Extended Data Figure 2. Validation of ESAM-based strategies to examine multipotent HSPC, erythropoiesis and B lymphopoiesis. a,** Overall experiment design (Top panels) and percentage of donor-derived cells (bottom panels) in lethally irradiated $CD45.1^+$ recipients transplanted with the indicated amount of $CD45.2^+$ SLAM (isolated as in Extended Data Fig. 1a) and ESAM (isolated as in Fig. 1e) LT-HSC (left) and ST-HSC (right) and $10^6$ $CD45.1^+$ total BM competitor cells (n = 7 recipient mice from 3 independent experiments per time point). Statistical analysis was performed using Two-way ANOVA, followed by Sidak's multiple comparisons test. It shows no significant difference for reconstitution ability between ESAM LT-HSC and SLAM LT-HSC, or ESAM ST-HSC and SLAM ST-HSC. **b,** Cell frequencies detected by FACS (white) or confocal imaging (orange) when using the strategy shown in Fig. 1e (n = 9 mice for each group). **c,** Representing images and quantification showing that ESAM correctly labels all CD31/CD144$^+$ vessels and CD41$^+$ megakaryocytes. (n = 3 sternum sections from 3 mice). **d,** CD150 and CD71 expression in indicated HSPC. **e,** FACS plots and colony-forming assays showing that Lin$^-$CD117$^+$Sca1$^-$CD41$^-$CD16/32$^-$CD105$^+$CD150$^+$ Pre CFU-E from Extended Data Fig. 1a are heterogeneous and contain CD71$^-$ and CD71$^+$ cells with the later fraction containing most erythroid colony-forming potential. The data also shows that CD71$^+$Ly6C$^+$ myeloid progenitors lack erythroid differentiation potential. (n = 3 in 3 independent experiments). **f,** FACS plots showing that mature hematopoietic cells do not coexpress CD117 and CD71 or CD117 and CD150. **g-j,** FACS gating strategies for isolation of -and representative images- of erythroid progenitors (g, h) terminal erythropoiesis (i, j). **k,** Frequencies for the indicated 8 populations in bone marrow by FACS (white) or image (orange) (n = 3 mice for each group). **l-m,** FACS gating strategies (l) for isolation of -and representative images (m) - of B lymphopoiesis (the Lin panel contains CD2, CD3e, CD5, CD8, CD11b, Ter119, Ly6G, IgM, and IgD). All data represent individual values with median plot. Statistical differences were calculated using two-tailed unpaired Student's t-test; *P* values are shown.

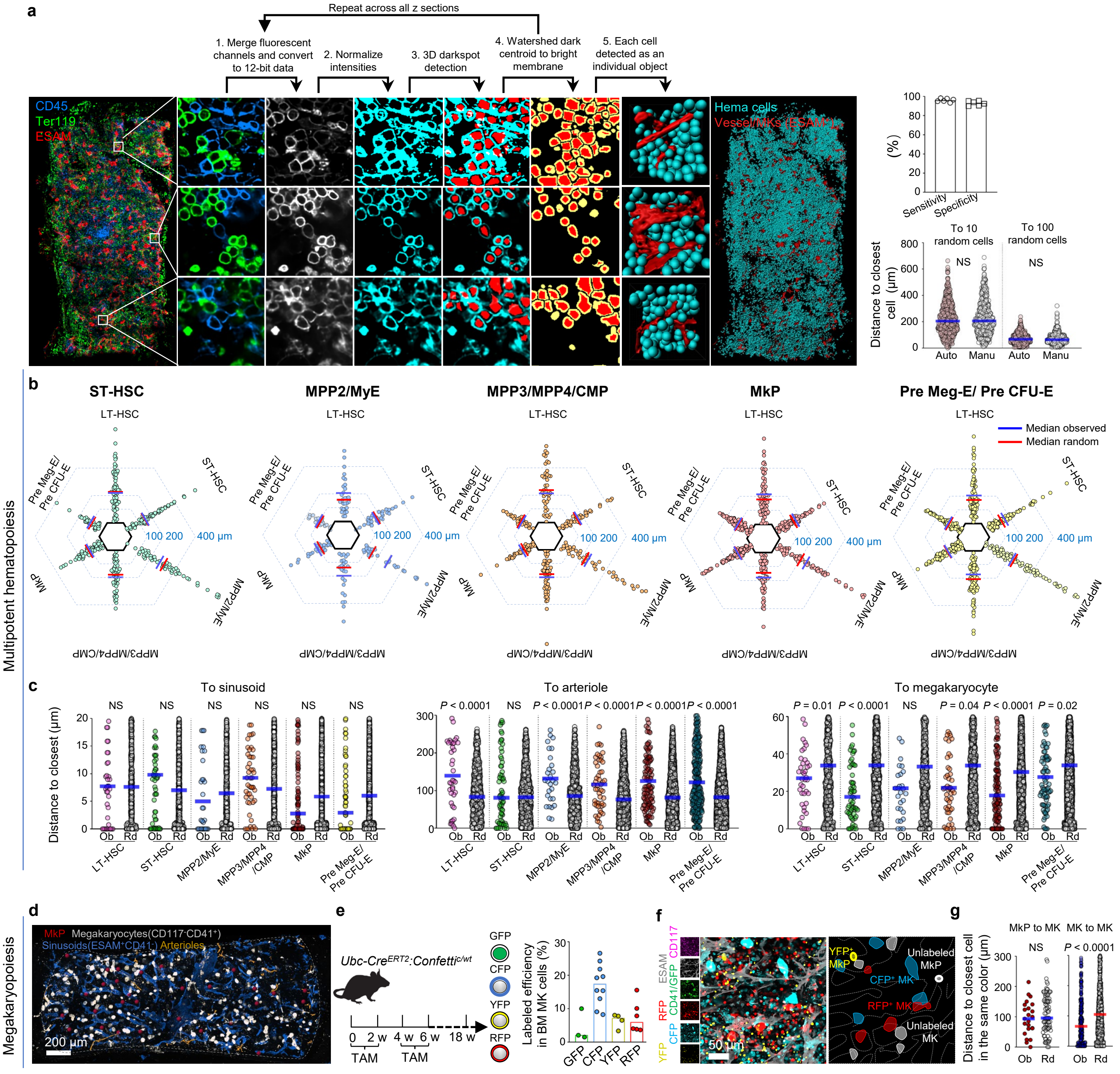

**Extended Data Figure 3. Mapping multipotent hematopoiesis and megakaryopoiesis in whole mounted sternum. a,** Experimental workflow for automatic cell segmentation. Fluorescence in the individual channels is merged, converted to 12-bit data, and intensities were normalized. The "3D darkspot detection" algorithm detects cells of different sizes. We then watershed each dark centroid to the bright membrane. This is repeated across all z sections until each cell is annotated as an individual object. The generated "inside cell" binary data was exported from Elements software to Imaris software and used to place dots representing each hematopoietic cell. The upper histogram shows sensitivity (= number of correctly segmented cells divided by the number of cells identified manually) and specificity (= number of correctly segmented cells divided by the number of identified cells using the automatic algorithm) of cell segmentation (n = 6 areas in 2 sternum segments from 2 mice). The lower histograms compare the distribution of distances between randomly selected cells segmented manually or using the automatic algorithm. **b,** Histograms showing the distribution of distances from each HSPC to the closest indicated cell (n = 66 ST-HSC, 30 MPP2/MyE, 41 MPP3/MPP4/CMP, 61 MkP, 82 Pre Meg-E/Pre CFU-E in 5 sternum segments from 4 mice). **c.** Distance analyses from each indicated cell to the closest sinusoid, arteriole, or megakaryocyte. (n = 42 LT-HSC, 63 ST-HSC, 32 MPP2/MyE, 55 MPP3/MPP4/CMP, 117 MkP, 85 Pre Meg-E in 5 sternum segments from 3 mice). **d,** Map showing the distribution of MkP and megakaryocyte (MK). **e,** Experimental design, and histogram showing the percentage of CFP, GFP, RFP, and YFP positive cells in bone marrow megakaryocytes in tamoxifen-treated *Ubc-cre*$^{ERT2}$*:confetti* mice. (Each dot represents one sternum segment from 3 confetti mice). **f,** Representative image showing confetti-labeled megakaryocyte progenitor (MkP) and megakaryocyte (MK) in tamoxifen-treated confetti mice. **g,** Observed (Ob) and Random cell (Rd) distances from each confetti-labeled MkP or MK to the closest MK labeled in the same color (n = 23 labeled MkP, 308 labeled MK in 3 sternum segments from 3 tamoxifen-treated confetti mice). Statistical differences were calculated using two-tailed unpaired Student's *t*-tests; *P* values are shown.

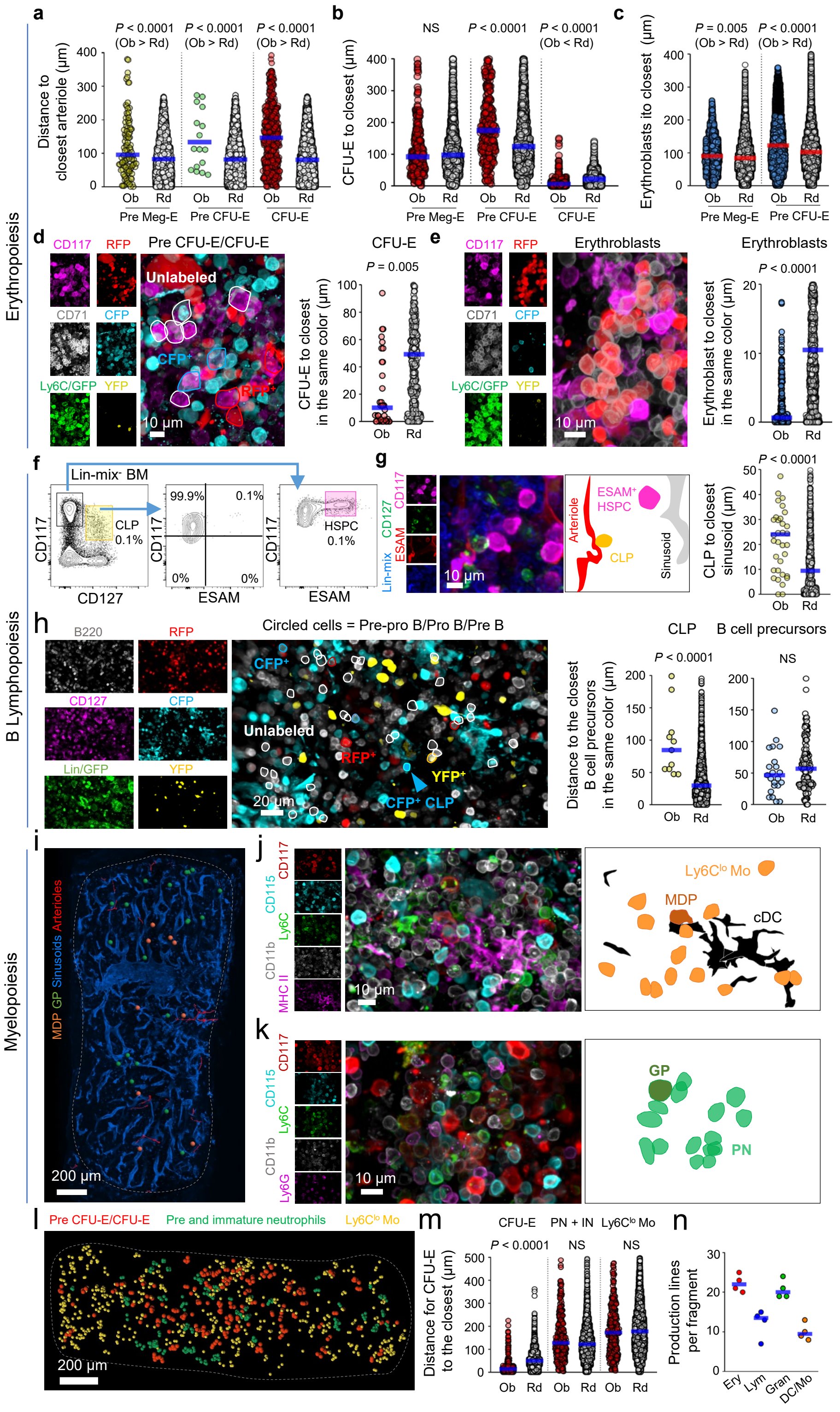


Extended Data Figure 4

**Extended Data Figure 4. Mapping erythropoiesis, B Lymphopoiesis and myelopoiesis in whole mounted sternum. a**, Distance analyses from each erythroid progenitor to the closest arteriole (n = 111 Pre Meg-E, 18 Pre CFU-E, and 523 CFU-E in 3 sternum segments from 3 mice). **b,** Distance analyses from CFU-E to the closest indicated cell (n = 318 CFU-E in 3 sternum segments from 3 mice). **c,** Distance analyses from erythroblasts ($CD117^{-}CD71^{+}Ly6C^{-}$, containing early and late erythroblasts, n = 4461 in 3 sternum segments from 3 mice) to the indicated erythroid progenitor. **d,** Representative images and distance analyses of confetti-labeled CFU-E to the closest CFU-E labeled in the same color (n = 38 confetti-labeled CFU-E in 5 sternum segments from 3 tamoxifen-treated confetti mice). **e,** Representative image showing confetti-labeled erythroblasts ($CD117^{-}CD71^{+}Ly6C^{-}$, containing early and late erythroblasts) in tamoxifen-treated confetti mice. Due to a lack of available fluorescence channels for analyses, Ly6C and GFP were combined in a single dump channel. The histogram shows distance analyses of each confetti-labeled erythroblast to the closest erythroblast labeled in the same color ($CD117^{-}CD71^{+}Ly6C^{-}$, containing early and late erythroblasts, n = 267 labeled erythroblasts in 4 sternum segments from tamoxifen-treated confetti mice). **f,** FACS gating strategy for simultaneous detection of $CD117^{+}CD127^{+}$Lin-mix$^{-}$CLP and $CD117^{+}ESAM^{+}$ HSPC (the Lin-mix panel contains B220, CD2, CD3e, CD5, CD8, CD11b, Ter119, Ly6G, CD41, Ly6C). **g,** Representative image and distance analyses showing that CLP map near arterioles and away from sinusoids. (n = 36 CLP in 3 sternum segments from 3 mice). **h,** Representative image and distance analyses of confetti-labeled CLP or B cell precursors ($B220^{+}Lin^{-}$ cells, the Lin panel contains CD2, CD3e, CD5, CD8, CD11b, Ter119, Ly6G, IgM, and IgD, containing all Pre-pro B, Pro B, and Pre B cells) to the closest B cell precursors labeled in the same color (n = 11 confetti-labeled CLP, 23 confetti-labelled B cell precursors in 3 sternum segments from 3 tamoxifen-treated confetti mice). **i,** Map showing monocyte dendritic progenitor cell (MDP) and granulocyte progenitor (GP) localization to sinusoids. **j, k,** Representative images showing a DC/$Ly6C^{lo}$ monocytes production line around an MDP (j), and a neutrophil production line centered around a GP (k). In all maps dots are three times the average size of the relevant cell. **l,** Map showing the distribution of Pre CFU-E / CFU-E, Pre and immature neutrophils, and $Ly6C^{lo}$ Monocytes ($Ly6C^{lo}$ Mo) in the sternum. Map dots are two times the average size of the relevant cells. **m,** Distance analyses from each CFU-E to the closest indicated cells (PN + IN: Pre and immature neutrophils) (n = 297 CFU-E in 3 sternum segments from 3 mice). n, Quantification of each type of production line per sternum segment (n = 4 sternum segments from 4 mice). Statistical differences were calculated using two-tailed unpaired Student's t-tests; *P* values are shown.

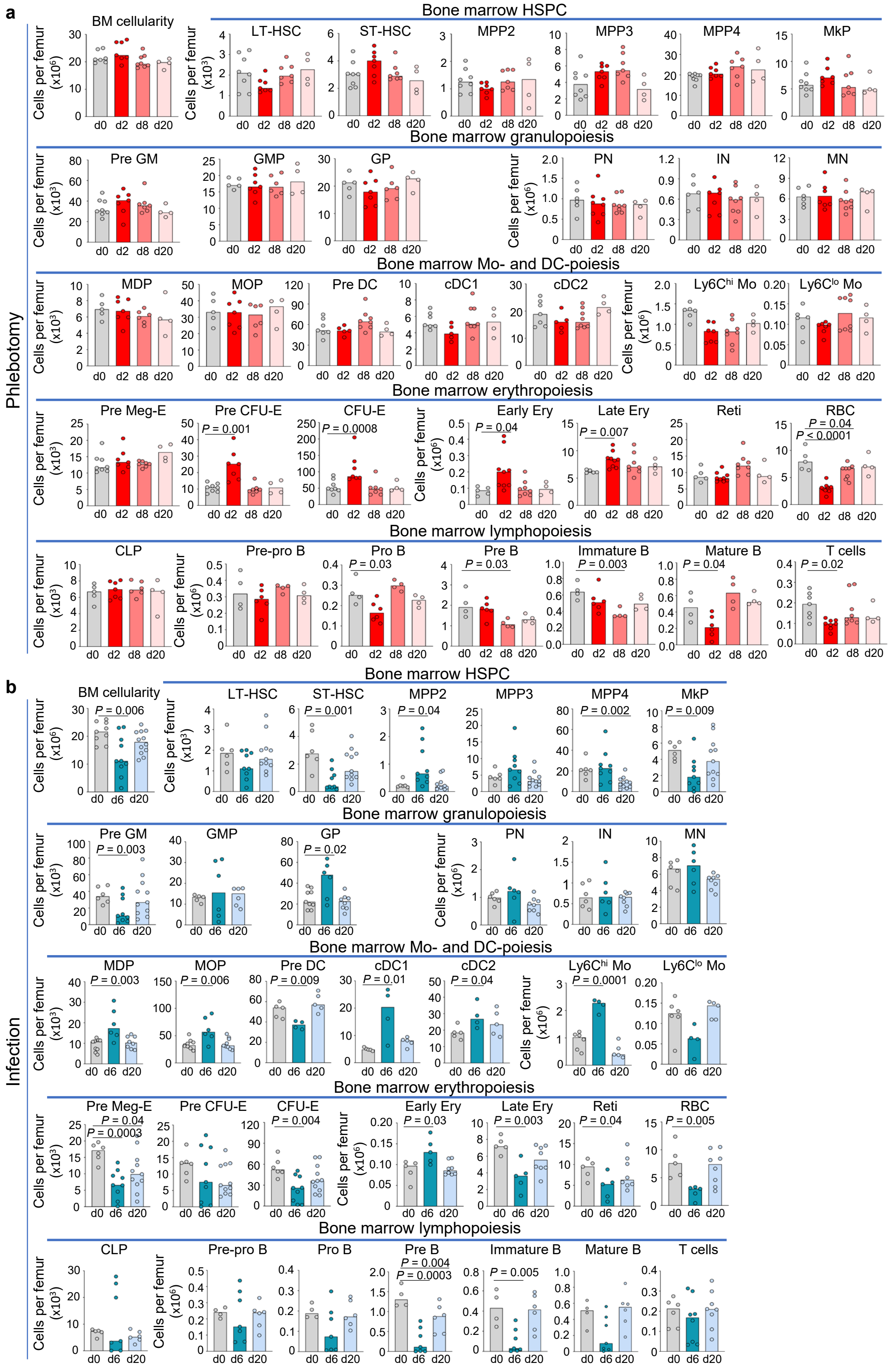


**Extended Data Figure 5. a, b**, Number of the indicated cells per femur (detected by FACS. Bone marrow HSPC populations are identified as in Extended Date Figure 1.a; granulopoiesis and Mo- and DC-poiesis populations are identified as described in reference[1]; erythropoiesis populations are identified as described in Extended Date Figure 2.g and i; lymphopoiesis populations are identified as described as in Extended Date Figure 2.l) at the indicated time points after phlebotomy (a) or *Listeria monocytogenes infection* (b). Each dot corresponds to one mouse in 3-6 independent experiments. (n = 4-12 mice for each time point). All data represent individual values with median plot. Statistical differences were calculated using two-tailed unpaired Student's *t*-tests; *P* values are shown.

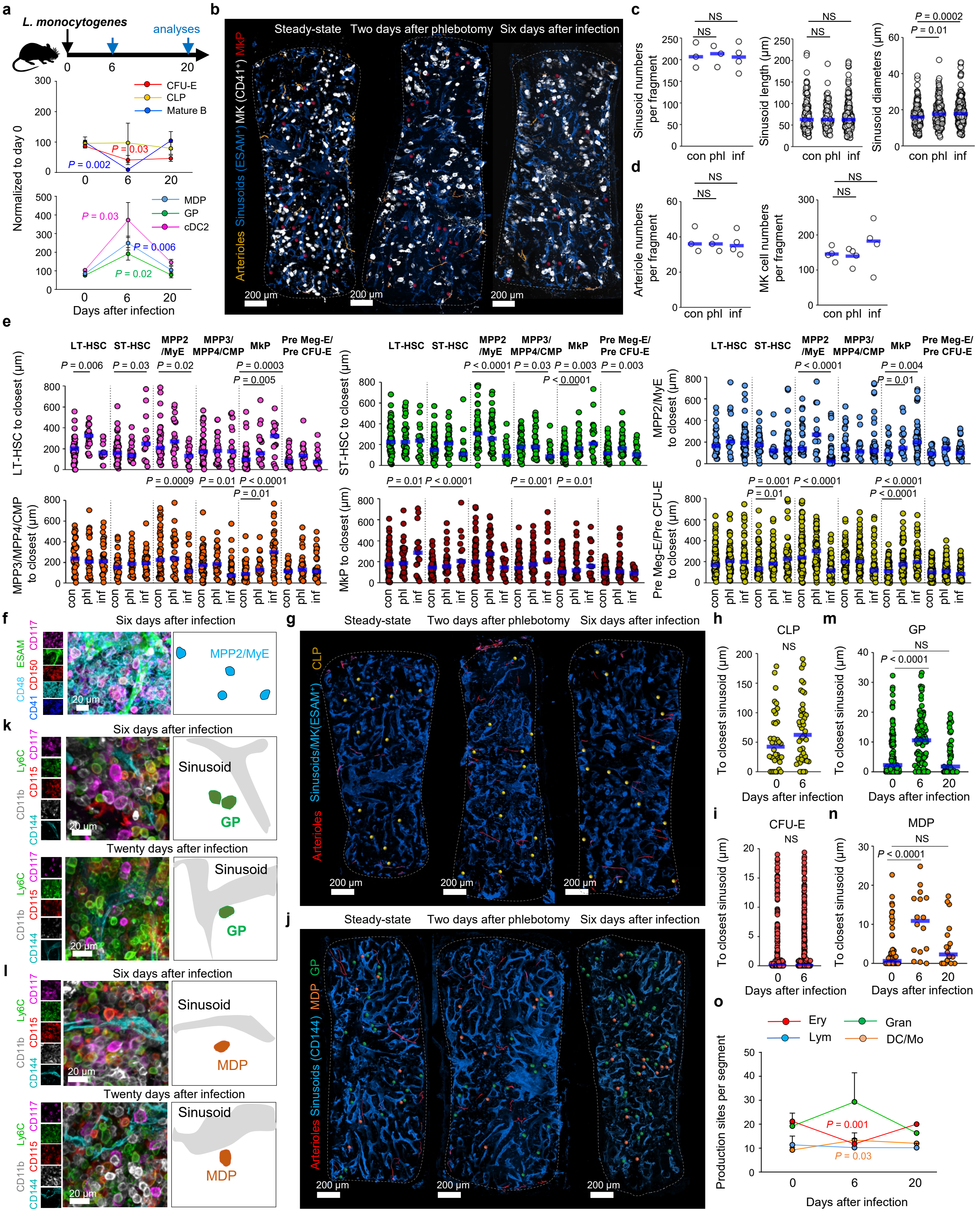


Extended Data Figure 6

**Extended Data Figure 6. The basic anatomy of hematopoiesis is largely maintained during acute stress. a,** Scheme showing the overall experiment design (Top panels) and quantification (bottom panels) of the indicated populations (normalized to day 0) in response to *L. monocytogenes* infection (n = 4 mice for each time point). All data represent mean $\pm$ s.e.m. **b-d,** Vascular organization in control (con), phlebotomized (phl) and *L. monocytogenes*-infected (inf) mice. Map (b); vessels numbers (n = 3, control and phlebotomy, or n = 4, infection, sternum segments from 3 mice), and sinusoid length and diameter (c, n = 50 randomly selected sinusoids from 3 sternum segments from 3 mice); arteriole and megakaryocyte numbers (d, n = 4, control, phlebotomy, or infection sternum segments from 4 mice). **e,** Distance analyses from each HSPC to the closest indicated cells at the indicated time points after challenge. (n = 41 LT-HSC,66 ST-HSC, 30 MPP2/MyE, 41 MPP3/MPP4/CMP, 61 MkP, and 82 Pre Meg-E in 4 sternum segments from 4 mice in steady-state for control; n = 21 LT-HSC, 35 ST-HSC, 16 MPP2/MyE, 30 MPP3/MPP4/CMP, 38 MkP, and 73 Pre Meg-E in 3 sternum segments from 3 mice two days after phlebotomy; n = 15 LT-HSC, 19 ST-HSC, 56 MPP2/MyE, 39 MPP3/MPP4/CMP, 17 MkP, and 57 Pre Meg-E in 3 sternum segments from 3 mice six days after infection). **f,** Representative image showing a loose MPP2/MyE cluster**. g-i,** Maps (g) and distance analyses showing CLP localization to sinusoid (h, steady-state (d0) n = 36, and six days after infection (d6) n = 43 CLP in 3 sternum segments from 3 mice) and CFU-E localization to sinusoids (i, steady-state (d0) n = 315, and six days after infection (d6) n = 314 CFU-E in 3 sternum segments from 3 mice). **j,** Maps showing MDP and GP distribution in the sternum after stress. **k-n**, Representative images (k, l) and quantification (m, n) showing transient GP and MDP detachment from sinusoids after infection. (n = 62, 16, 18 MDP; and n = 114, 98, 49 GP in 3-5 sternum segments from 3 mice in steady-state (d0), six days (d6), and twenty days (d20) after infection). **o,** Quantification of each type of production line per sternum after infection (n = 3-5 sternum segments from 3-4 mice for each time point). All data represent mean $\pm$ s.e.m. Dots in maps are three times the size of the relevant cell except for CLP dots which are five times the actual size. Statistical differences were calculated using two-tailed unpaired Student's *t*-tests; *P* values are shown.

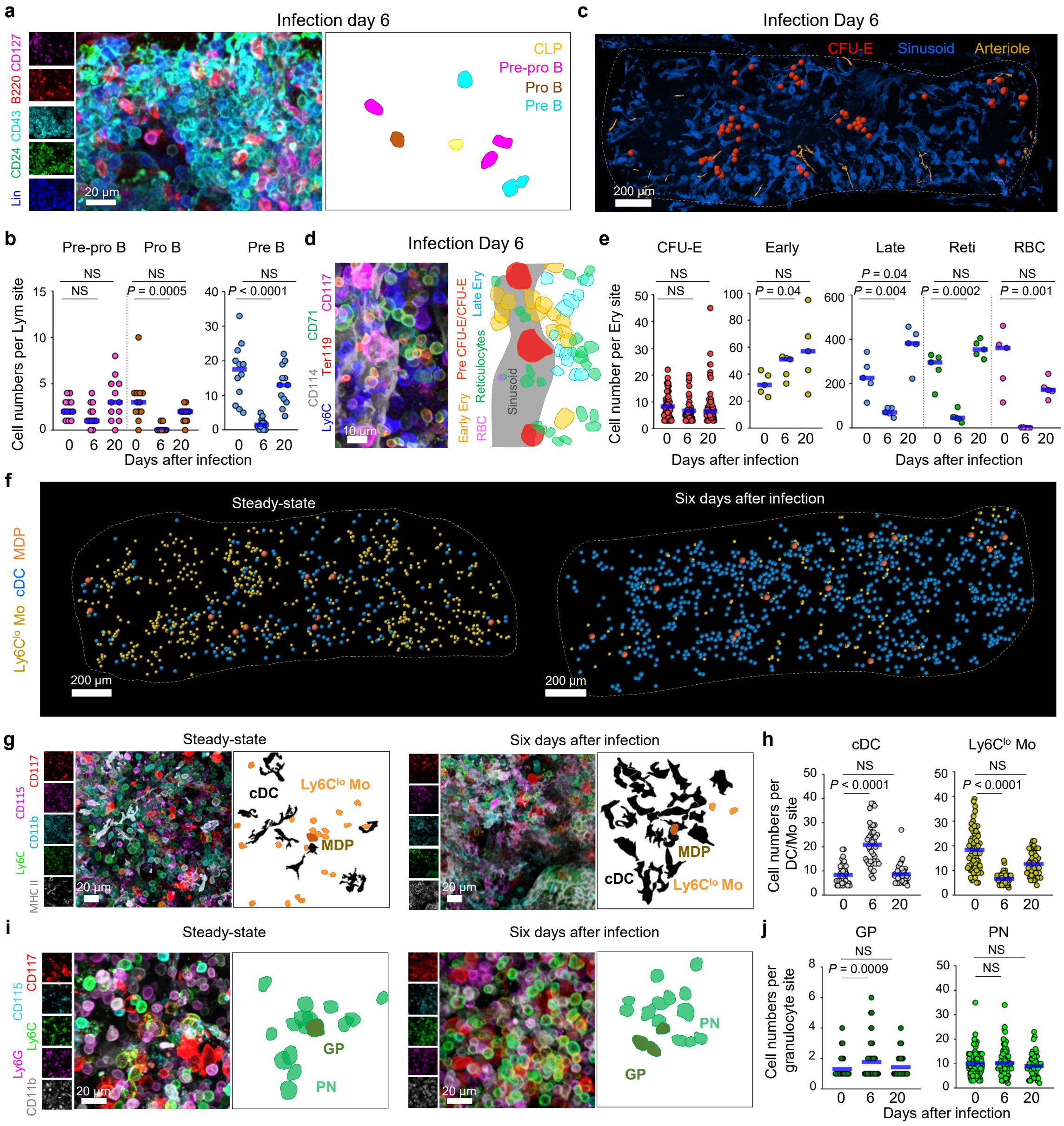

**Extended Data Figure 7. Production lines orchestrate hematopoietic plasticity to stress. a, b,** Representative images (a) and number of cells per B cell line (b) from control or *L. monocytogenes*-infected mice after infection (n = 12 randomly selected production lines in 3 sternum segments from 3 mice for each time point). **c,d,** Map and representative images showing the changes in erythroid production lines after infection. Map dots are three times the average size of the relevant cell. **e**, Number of CFU-E (n = 65, 45, and 60 production lines for days 0, 6, 20, three sternum segments from 3 mice per time point) and indicated erythroid cells (n = 5 randomly selected production lines for each time point, in 3 sternum segments from 3 mice) from control or *L. monocytogenes*-infected mice. **f-h,** Maps (f), Representative images (g) and indicated cell number (h) in DC/Ly6C$^{lo}$ monocyte production lines from control or *L. monocytogenes*-infected mice (n = 49, 35, and 29 randomly selected production lines for days 0, 6, 20, in 3-5 sternum segments from 3 mice per time point). **i, j,** as g-h but for granulocyte production lines (n = 96, 88, and 50 randomly selected production lines in 3-5 sternum segments from 3-4 mice per time point). Dots in maps are three times the size of the relevant cell. Statistical differences were calculated using two-tailed unpaired Student's *t*-tests; *P* values are shown.

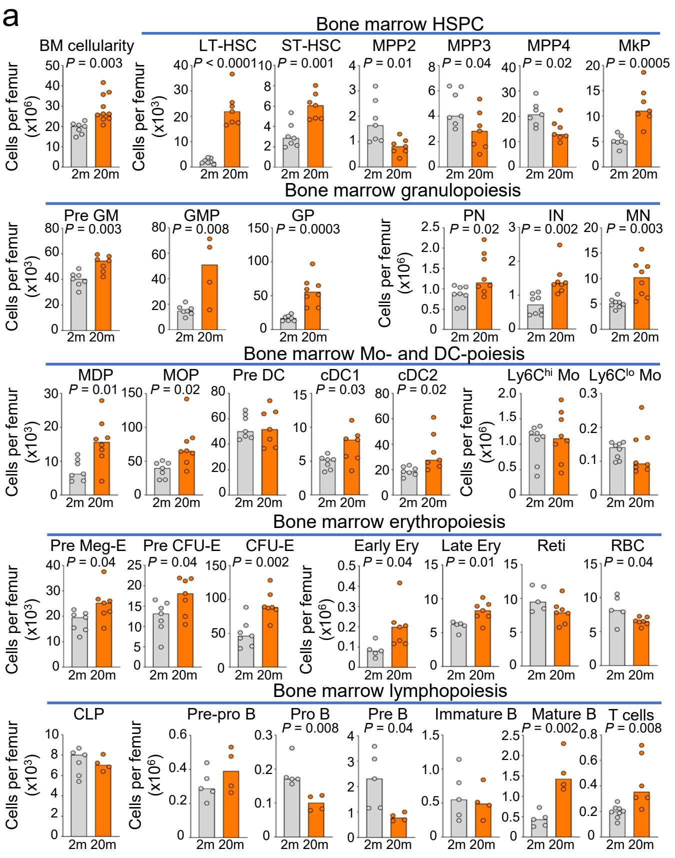


**Extended Data Figure 8. Quantification of bone marrow cell number changes in aged mice. a,** Number of BM cells or the indicated populations per femur (Bone marrow HSPC populations are identified as in Extended Date Figure 1.a; granulopoiesis and Mo- and DC-poiesis populations are identified as described in reference[1]; erythropoiesis populations are identified as described in Extended Date Figure 2.g and i; lymphopoiesis populations are identified as described as in Extended Date Figure 2.l) in 2-month-old (n = 7) and 20-month-old mice (n= 4-10). Each dot corresponds to one mouse in 3 independent experiments. All data represent individual values with median plot. Statistical differences were calculated using two-tailed unpaired Student's *t*-tests; *P* values are shown.

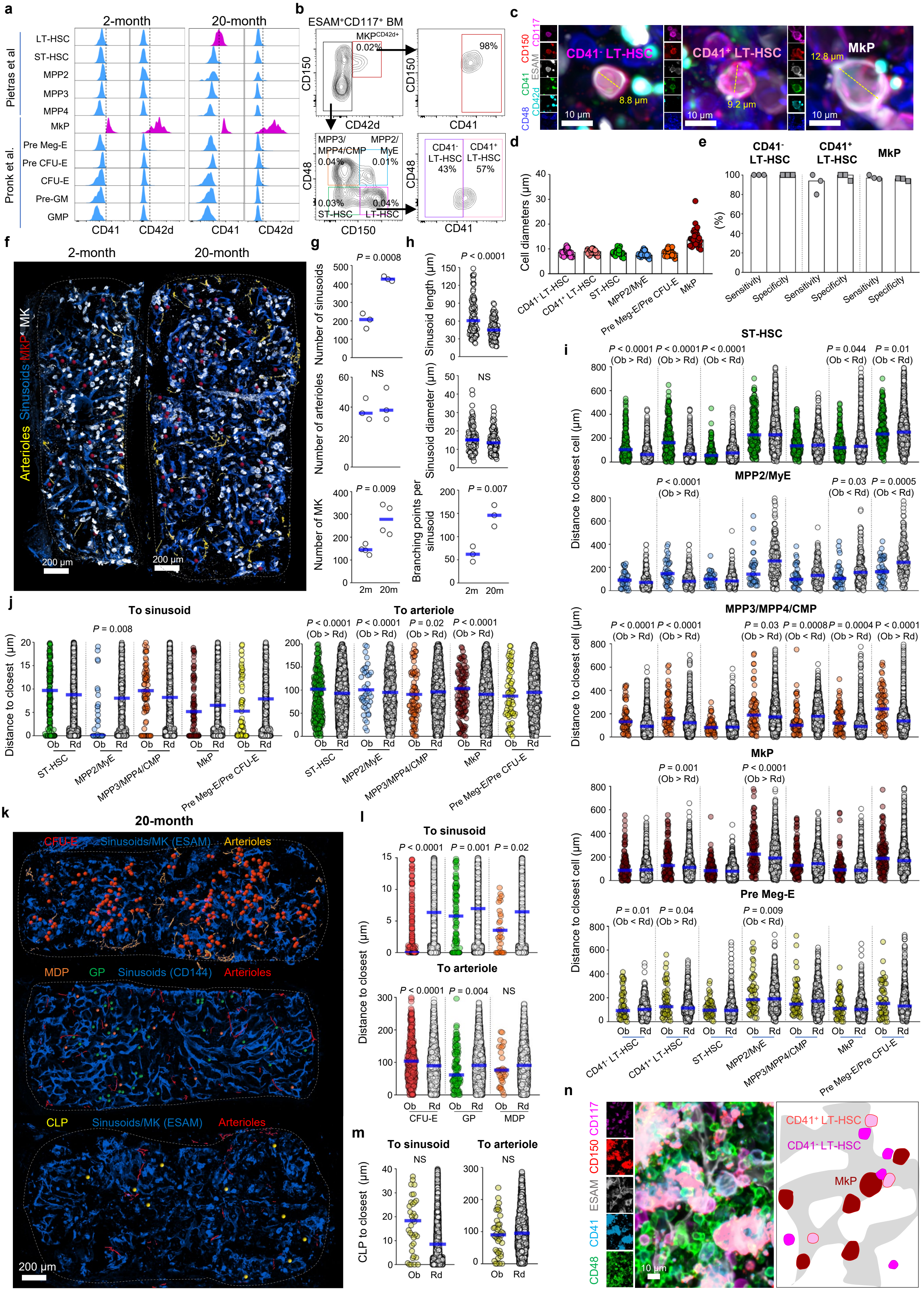


Extended Data Figure 9

**Extended Data Figure 9. The basic anatomy of hematopoiesis is maintained in aged mice. a,** Histograms showing CD41 and CD42d expression in the indicated HSPC in 2- and 20-month old mice. **b,** FACS plots showing the gating strategy to identify indicated HSPC with CD42d and CD41 expression in MkP and LT-HSC in 20-month-old mice. **c, d,** Representative images (c) and quantification of cell diameters (d) demonstrating that ESAM$^{+}$CD117$^{+}$CD48$^{-}$CD150$^{+}$CD42$^{-}$CD41$^{+}$ myeloid-biased LT-HSC, ESAM$^{+}$CD117$^{+}$CD48$^{-}$CD150$^{+}$CD42$^{-}$CD41$^{-}$ lymphoid-biased LT-HSC and ESAM$^{+}$CD117$^{+}$CD150$^{+}$CD42$^{+}$CD41$^{+}$ MkP can be distinguished based on cell size (n = 30 randomly selected cells from each type in 3 sternum segments from three 20-month-old mice). **e,** Sensitivity (= Number of ESAM$^{+}$CD117$^{+}$CD48$^{-}$CD150$^{+}$CD41$^{-}$ LT-HSC, ESAM$^{+}$CD117$^{+}$CD48$^{-}$CD150$^{+}$CD41$^{+}$ LT-HSC, ESAM$^{+}$CD117$^{+}$CD150$^{+}$CD41$^{+}$ MkP correctly identified based on CD150 and CD41 expression and cell size, divided by the number of cells identified when counterstained with CD42 (CD41$^{-}$CD42$^{-}$ LT-HSC, CD41$^{+}$CD42$^{-}$ LT HSC, and CD41$^{+}$CD42$^{+}$ MkP) and specificity (= Number of ESAM$^{+}$CD117$^{+}$CD48$^{-}$CD150$^{+}$CD41$^{-}$ LT-HSC, ESAM$^{+}$CD117$^{+}$CD48$^{-}$CD150$^{+}$CD41$^{+}$ LT-HSC, ESAM$^{+}$CD117$^{+}$CD150$^{+}$CD41$^{+}$ MkP correctly identified based on CD150 and CD41 expression and cell size, divided by the number of cells identified based on CD150 and CD41 expression and cell size) for distinguishing myeloid and lymphoid biased LT-HSC and MkP based on CD41 and CD150 expression and cell size (compared with CD42d based identification, 47 CD41$^{+}$ LT-HSC, 55 CD41$^{-}$ LT-HSC, and 70 MkP in n = 3 sternum segments from two 20-month-old mice were analyzed). **f-h,** Age-dependent changes in the vascular microenvironment. Maps (f), number of sinusoids, arterioles, and megakaryocytes (g, n = 3 sternum segments from three mice for vessel quantification and n = 4 sternum segments from four 2- and 20-month-old mice for megakaryocyte quantification); and sinusoid length, diameter, and branching points (h, n = 50 sinusoids randomly selected in 2 sternum segments from two 2- and 20-month-old mice). **i,** Distance analyses from each HSPC to the closest indicated cells in 20-month-old mice. (n = 191 CD41$^{-}$ LT-HSC, 109 CD41$^{+}$ LT-HSC, 236 ST-HSC, 39 MPP2/MyE, 72 MPP3/MPP4/CMP, 133 MkP and 57 Pre Meg-E in 5 sternum segments from three 20-month-old mice). **j**, Distance analyses from each HSPC to the closest sinusoid and arteriole. (n = 315 ST-HSC, 46 MPP2/MyE, 102 MPP3/MPP4/CMP, 131 MkP and 79 Pre Meg-E in 5 sternum segments from three 20-month-old mice). **k**, Maps showing committed progenitor distribution in the aged marrow. **l**, Distance analyses from each committed progenitor to the closest sinusoid or arteriole (n = 558 CFU-E, 115 GP, 33 MDP in 3 sternum segments from three 20-month-old mice). **m**, Distance analyses showing a lack of CLP localization to arterioles or sinusoids in 20-month-old mice (n=38 CLP in three sternum segments from three 2- and 20-month-old mice) in aged mice. Dots in maps are 3-times (CFU-E, GP, and MDP), or 5-times (CLP) the size of the relevant cell. **n,** Representative images showing an LT-HSC cluster in the aged sternum. Statistical differences were calculated using two-tailed unpaired Student's *t*-tests; *P* values are shown.

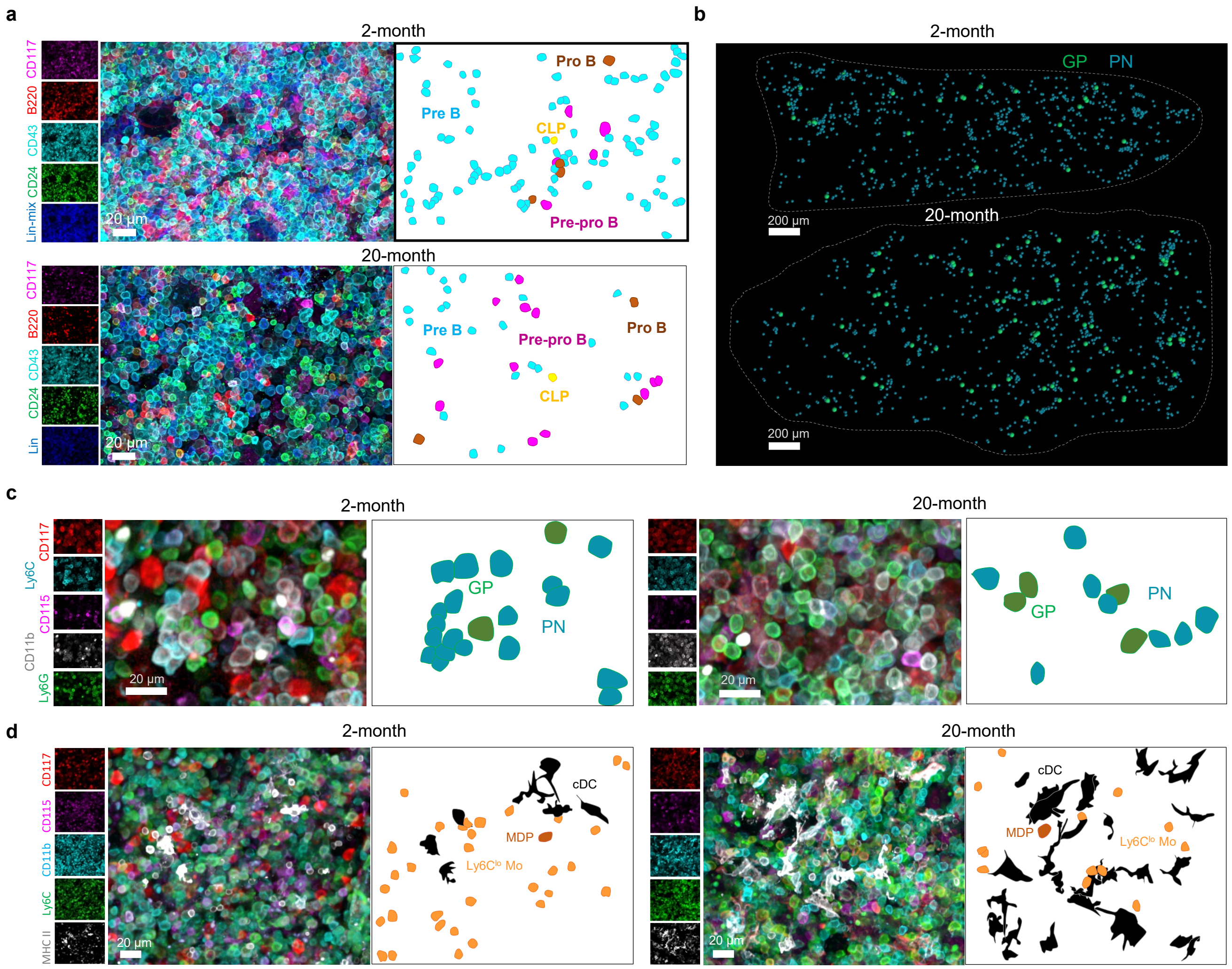


**Extended Data Figure 10. Aging caused extensive cell and lineage-specific microanatomical adaptations in each type of production line. a,** Representative images showing stage-specific delayed B cell differentiation (the Lin panel contains CD2, CD3, CD5, CD8, CD11b, Ter119, Ly6G, IgM, IgD). **b-d,** Maps (b) and representative images showing delayed differentiation in granulocyte production lines (c) and increased dendritic cell and reduced monocyte output (d) in dendritic cell (DC)/Ly6C$^{lo}$ monocytes (Ly6C$^{lo}$ Mo) production lines in aged mice. Dots in maps are 3-times (GP), or 2-times (PN) the size of the relevant cell.
